# Fibrin-containing Hydrogels Regulate Human Astrocyte State and Neuronal Reprogramming

**DOI:** 10.1101/2025.11.05.686711

**Authors:** Thomas Distler, Katherina Konrad Daga, Martina Bürkle, Sebastián Vásquez Sepúlveda, Kristian Franze, Magdalena Götz, Giacomo Masserdotti

## Abstract

Astrocytes are key components in reactive gliosis after brain injury, yet defined *in vitro* models dissecting the influence of extracellular matrix (ECM) components enriched after injury, such as fibrin, on human astrocyte behaviour and function are still missing. Here, we use fibrinogen-derived fibrin and fibrin-alginate-RGD (FAR) 3D hydrogel substrates to examine the influence on human iPSC-derived astrocyte behaviour and their direct conversion into neurons. Astrocytes develop complex morphologies in 3D-FAR hydrogels while are more proliferative and migratory in 3D-Fibrin. Interestingly, gene expression profile analysis revealed different reactive states of astrocytes in 3D-Fibrin and 3D-FAR, which persist over time. The highly inflammatory state in 3D-FAR is largely incompatible with direct neuronal reprogramming hampering the direct conversion even at early stages. Conversely, astrocytes in 3D-Fibrin hydrogels can readily convert into neurons, demonstrating a potent influence of how fibrin is presented on eliciting distinct astrocyte states with great relevance for fate conversion.

**Research highlights:**

- First transcriptome of human astrocytes in 3D-Fibrin hydrogel and derivative
- Fibrin-alginate-RGD (3D-FAR) hydrogel elicits high branching complexity along with exacerbated reactive signature in astrocytes
- 3D-Fibrin hydrogels enable proliferation and migration of astrocytes
- Direct conversion of human iPSC-derived astrocytes into neurons in 3D-Fibrin hydrogel

## Introduction

Astrocytes are the most abundant glial cell-type in the mammalian central nervous system (CNS) ^1,2^ and play key roles in pathological conditions ^3–5^. As such, they represent a promising target for therapeutic strategies by fostering their beneficial effects and reducing their adverse functions ^6–8^: for example, the increased proliferation of juxtavascular astrocytes reduces the invasion of monocytes and scar formation ^9–11^, while neurotoxic astrocytes hamper neuron function and survival ^12–14^.

Blood-derived fibrinogen, and its polymerised product fibrin, directly affect border-forming and proliferating astrocytes in mouse models ^15^ but also regulate the expression and release of extracellular matrix (ECM) proteins ^15^: this dual effect complicates investigating fibrinogen specific effects from those of other components within the complex *in vivo* environment. Moreover, still very little is known about its influence on human astrocytes ^8^, besides the general use of fibrin as substrate in blood brain barrier models, which support the growth of stem cell-derived human astrocytes ^16,17^. Therefore, we set-out to examine the impact of fibrin-rich 3D environments on human induced pluripotent stem cell (iPSC)-derived astrocytes ^18,19^ as fibrin is a key driver of lesion border-forming astrocytes^15^, and generate 3D fibrin hydrogels (hereafter, 3D-Fibrin), which reflect the soft nature of brain tissue ^20^ after acute injury versus a combination of fibrin and RGD-peptide coupled alginate (hereafter, 3D-FAR), which shows higher stiffness. This allowed to identify profound differences in astrocyte states in 3D soft and stiffened environments, with remarkably potent influence on their capacity to be directly reprogrammed into neurons.

Even though proliferating astrocyte populations emerge after injury ^20^, they fail to replace neurons which are lost after injury. However, proliferating astrocytes partially dedifferentiate assuming hallmarks of neural stem cells (NSCs), whose neurogenic capacity can be revealed *in vitro,* after removing them from the *in vivo* gliogenic environment ^11^, or by further manipulations, such as Notch deletion ^21^ or the expression of neurogenic fate determinants ^22^. Excitingly, such proliferating astrocyte population was recently discovered in human pathologies ^23^. However, little is known about which factors may impede the conversion of these in principle plastic astrocytes after brain injury. To tackle this question, we used the recently established reprogramming paradigm of human iPSC-derived astrocytes into neurons^19^, and applied it to different 3D hydrogels (3D-FAR or 3D-Fibrin). Interestingly, the former virtually completely interfered with the direct conversion into neurons, correlating to a highly inflammatory reactive state, as assessed by gene expression profile. Conversely, 3D-Fibrin gels allowed astrocytes to convert into neurons, showing the relevance and importance of modelling individual and combined ECM components in 3D for human astrocyte states and their response to fate conversion.

## Results

### Human astrocytes acquire branching in 3D-FAR hydrogels

To test if 3D environment influences human iPSC-derived astrocytes ^18,19,23^, we first expanded the cells for 35 days in EGF/FGF-containing astrocyte medium (AM) and then embedded them in four different 3D hydrogels (Figures 1A and S1A): a) soft sodium alginate at low concentration and molecular weight (0.5% w/v, 40kDa), pre-functionalized with RGD peptide (GRGDSP, degree of substitution (DOS): 0.375%, from manufacturer) as integrin ligands ^24^ at low density; b) soft fibrin, offering different RGD sites by fibrinogen polymerisation (e.g. Aα95–98 (RGDF) ^25,26^, Aα572–575 (RGDS) ^25,26^, bovine: Aα252-254 ^27^); c) Cultrex^TM^, derived from basement membrane (BME), and d) fibrin-<u>a</u>lginate-<u>R</u>GD hydrogels (FAR), providing adhesion sites through both fibrin and alginate-RGD (Figure S1A). Seven days after seeding or embedding (days post embedding, dpe), astrocytes on 2D-PLO-laminin-coated glass coverslips (hereafter, 2D-POL), used as control, were flat (Figure S1A) and cells in alginate-RGD remained mainly round. Remarkably, in 3D-Fibrin and Cultrex^TM^ astrocytes showed an elongated morphology and network-like structure (Figure S1A), while they were more ramified in 3D-FAR hydrogels. This observation suggested that the fibrin and alginate-RGD in 3D-FAR have a synergistic rather than an additive effect. Therefore, we focused on the behaviour of astrocytes embedded either in 3D-Fibrin or in 3D-FAR hydrogels.

**Figure 1.**
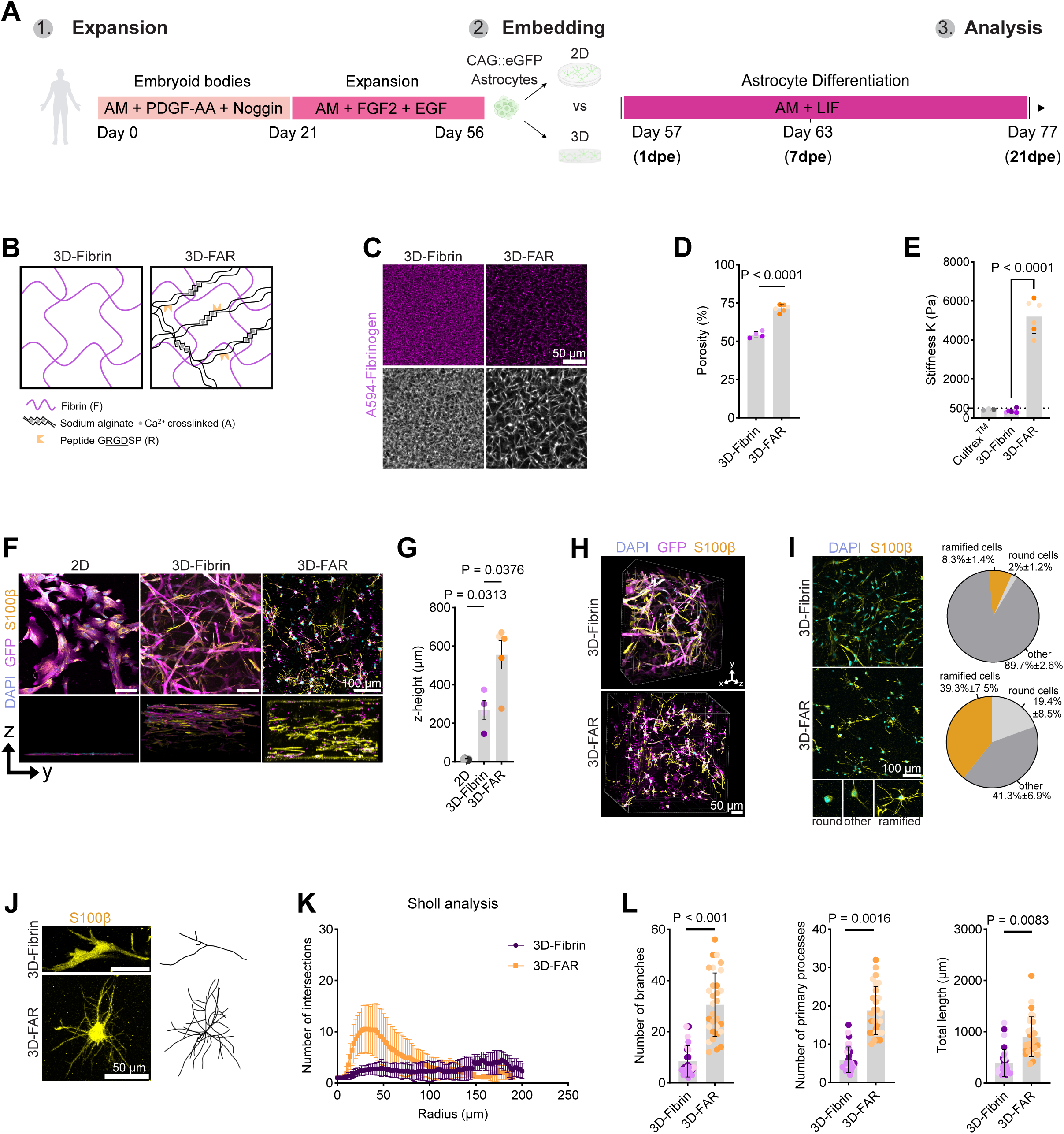
Characterization of human iPSC-derived astrocytes in 3D hydrogels. (**A**) Experimental design: iPSC-to-astrocyte differentiation protocol (left panel); embedding of astrocytes in 3D hydrogels or seeded on 2D-POL-coated coverslips (right panel). (**B**) Schematic illustration of 3D-Fibrin and 3D-FAR hydrogels. (**C**) Confocal microscopy images of 3D-Fibrin and 3D-FAR microstructure by AF594 labelling of fibrinogen. (**D**) Graph showing the percentage of microporosity, quantified as the volume between polymerized fibrin fibres in segmented images (N = 3; two technical gel replicates per N for fibrin and FAR; two-tailed t-test, *p < 0.0001*). (**E**) Graph showing the apparent elastic modulus (stiffness) K_eff_ of 3D-Fibrin, 3D-FAR hydrogels and Cultrex^TM^ (N = 3 independent preparations and two hydrogel replicates per preparation [one technical replicate in case of Cultrex^TM^]; two-tailed t-test, *p < 0.0001*). (**F**) Examples of astrocytes in 2D-POL-coated coverslips or in 3D-Fibrin and 3D-FAR hydrogels at 7 days post embedding (dpe). (**G**) Graphs showing thickness (z-height) of hydrogels containing cells (N ≥ 4; mean ± S.E.M; one-way Anova, *p < 0.05*). (**H**) 3D image of astrocytes embedded in 3D-Fibrin and 3D-FAR at 7dpe. (**I**) Pictures (left panels) showing astrocytes morphology embedded in 3D-Fibrin or 3D-FAR: pie charts (right panel) showing the proportion of ramified astrocytes (S100β^+^ cells with ≥ three main processes) inside 3D-Fibrin and 3D-FAR hydrogels at 7 dpe (N=3 biological replicates; n=10 cells analysed per condition per biological replicate). (**J**) Examples of astrocyte morphologies in 3D-Fibrin or 3D-FAR hydrogels at 7 dpe. (**K, L**) Sholl analysis and morphology characterisation of S100β astrocytes grown in 3D-Fibrin or 3D-FAR hydrogels at 7 dpe (n = 30 astrocytes per condition, from N = 3 biological experiments; different colours represent astrocytes from the same biological experiment. Data are shown as mean ± SD. Statistical analysis was performed on averages from each biological experiment using students t-test).

To get insights into the properties of the two hydrogels, we analysed their microstructure and stiffness, potentially influenced by the presence of Ca^2+^ ions used to crosslink alginate (Figure 1B). Performing multiphoton-microscopy of 3D-Fibrin and 3D-FAR hydrogels containing AF594-conjugated fibrinogen revealed an increased porosity between fibrin-fibres in 3D-FAR (Figures 1C and D) and different microstructure (Figure 1C), likely due to the presence of the alginate matrix. To assess the mechanical properties of the hydrogels, we performed atomic force microscopy (AFM) of 3D-Fibrin and 3D-FAR (Figure 1E) and included also Cultrex^TM^ ^28,29^, previously used for mouse astrocytes in 3D ^30^. 3D-FAR hydrogels were 10x stiffer (K_eff_ ∼5204 Pa, p < 0.0001) than 3D-Fibrin (K_eff_ ∼352 Pa), or Cultrex^TM^ BME gels (K_eff_ ∼438 Pa), which are in the range of neonate-to-adult murine brain stiffness (∼240 – 480 Pa ^31^) and human gliotic tissue^32^. The stiffness of 3D-FAR hydrogels was in the range of human WHO grade IV glioblastoma brain tissue (∼3 kPa - 13.5 kPa^32^) and in accordance to previous observations that CaCl_2_ can tune alginate stiffness^33,34^ and reinforce other hydrogel matrices^35^. Importantly, the stiffness measured in 3D-Fibrin and 3D-FAR gels was reproducible across different fibrinogen batches (Figure S1B).

Given the mechanical differences between 3D-Fibrin and 3D-FAR hydrogels, we examined the behaviour of GFP-labelled astrocytes by continuous live imaging after embedding them either in AF594-conjugated 3D-Fibrin (Video S1) or AF594-conjugated 3D-FAR (Video S2) (Figures S1C-F). While astrocytes could actively remodel 3D-Fibrin hydrogel (Video S1), and navigate through it (Video S3), 3D-FAR-embedded astrocytes were relatively immobile, rather extending and remodelling their processes (Video S2). We also observed cell division followed by cell death in 3D-FAR hydrogels (Video S4).

Overall, astrocytes cultured in both 3D substrates spread three-dimensionally (over ∼250 µm of thickness in 3D-Fibrin, up to ∼500 µm in 3D-FAR; Figures 1F-1H, Video S5 - S7). Morphological analysis at 7 dpe supported the observations from continuous live imaging: ∼39% of S100β^+^ astrocytes inside 3D-FAR showed complex morphologies (equal or more than three primary branches), while the rest were either less branched (∼41%), or round (∼19%) (Figure 1I). Conversely, in 3D-Fibrin a small proportion of S100β^+^ astrocytes showed a complex morphology (∼8%), with almost 90% of the cells showing a simple (1-2 primary branches) or round (∼2%) morphology. Noteworthy, Sholl analysis (Figures 1J-L) revealed that ramified astrocytes in 3D-FAR are significantly more complex than 3D-Fibrin-embedded counterparts (primary branches: 3D-FAR 20 ± 6, 3D-Fibrin 8 ± 2, *p* = 0.0016; total number of branches: 3D-FAR 31 ± 11, 3D-Fibrin 7 ± 2, p < 0.001; Figure 1L) and overall had longer processes (summed branch length: 3D-FAR ∼900 µm, 3D-Fibrin ∼388 µm, *p* = 0.0083; Figures 1J-L). Together, the data revealed that iPSC-derived astrocytes can be embedded in 3D hydrogels with different mechanical properties and microstructure, and this strongly affects the engagement of the cells with the substrates and their morphology.

### Analysis of cell viability and proliferation of astrocytes in 3D-Fibrin and 3D-FAR

At 7 dpe more than 90% of GFP-labelled astrocytes in both 3D environments were S100β positive, but a higher proportion of cells were positive for the astroglial marker glial fibrillary acidic protein (GFAP) in 3D-FAR, a marker usually upregulated upon reactive gliosis ^12^, (Figures 2A and 2B), thus suggesting a more reactive state in 3D-FAR hydrogels than in 3D-Fibrin (see below).

**Figure 2.**
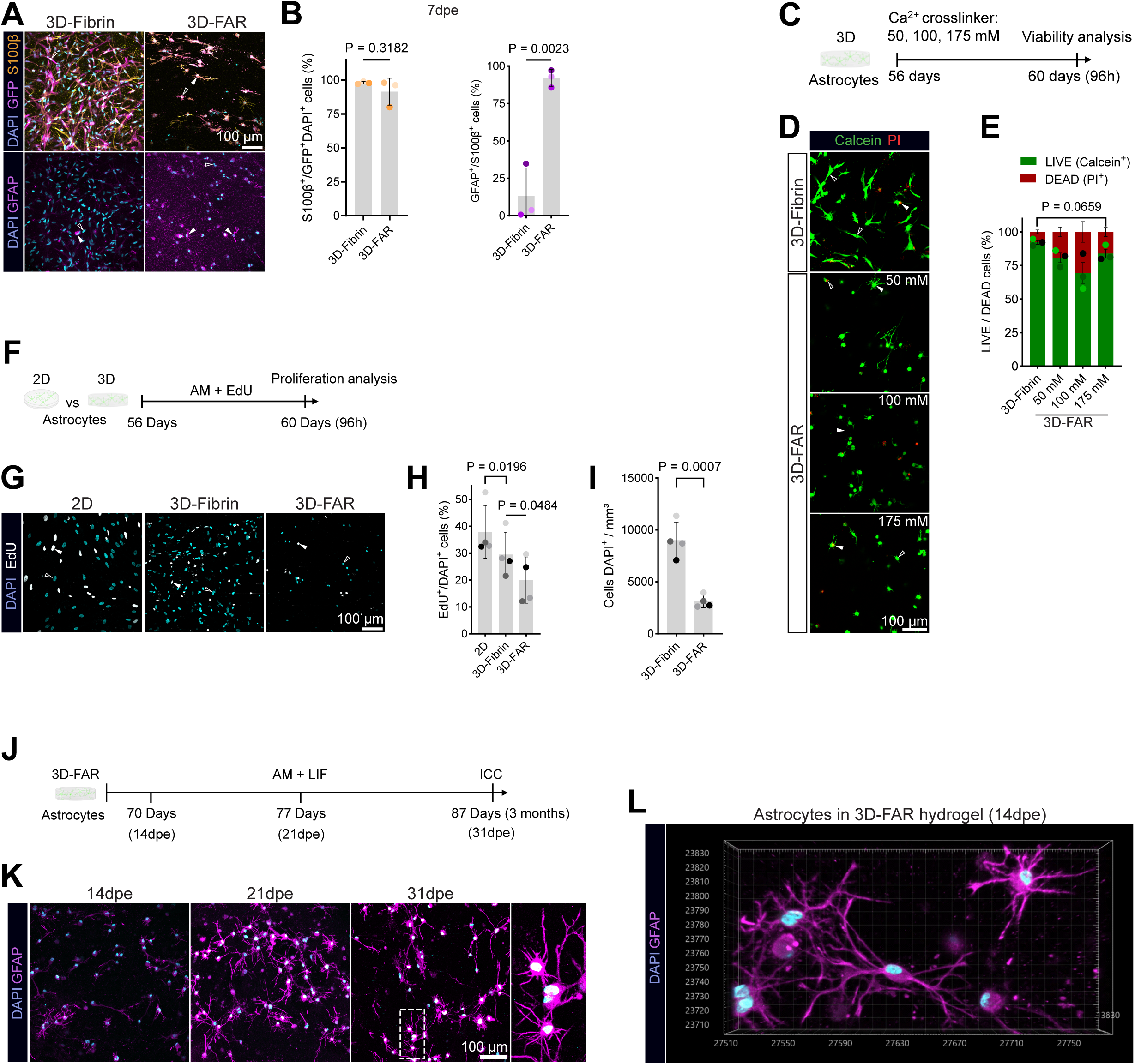
Analysis of astrocyte phenotypes in Fibrin and 3D-FAR hydrogels. **(A)** Confocal microscopy images of S100β^+^ and GFAP^+^ astrocytes in 3D-Fibrin or 3D-FAR hydrogels at 7dpe. Scalebar = 100µm. **(B)** Graphs showing the proportion of S100β^+^ (left panel) and GFAP^+^ (right panel) cells in 3D-Fibrin and 3D-FAR hydrogels (N = 3; unpaired students t-test. Data shown as mean ± SD). **(C)** Scheme of viability assay in astrocytes following the use of different concentration of calcium chloride (50, 100, 175 mM) for crosslinking 3D-FAR hydrogel. **(D)** Examples of astrocytes positive for Calcein^+^ (alive cells, green), propidium iodide (PI^+^, dead cells, red). Scalebar = 100µm. **(E)** Graph showing the proportion of alive or dead cells after 4 days in either 3D-Fibrin or 3D-FAR crosslinked with 0.175 M CaCl_2_ (N = 4; mean ± S.E.M; Data point colours indicate different biological replicates. Paired students t-test). **(F, G)** Scheme (**F**) and pictures (**G**) of EdU-incorporation assay to assess proliferation of astrocytes in 3D-Fibrin, 3D-FAR, and on 2D-POL glass, after 96h. Scalebar = 100µm. **(H, I)** Graphs showing the proportion of EdU^+^ cells (**H**) and cell density (**I**) in 3D-Fibrin and 3D-FAR gels (N = 4; mean ± SD. Paired two-tailed students t-test.) **(J-L)** Scheme (**J**) and pictures of GFAP^+^ astrocytes (**K**) cultured in 3D-FAR for different time points. Astrocytes form cell-cell contacts by 14 dpe (**L**). Scalebar = 100µm.

Next, we evaluated whether the concentration of Ca^2+^ (175mM) used to crosslink alginate in 3D-FAR could contribute to the cell death observed in live-imaging and, thus, explain the lower cell density found in this substrate (Video S5 and S6). To this end, we crosslinked alginate-RGD inside 3D-FAR hydrogels with different Ca^2+^ concentrations (50mM, 100mM and 175mM) and, four days later, assessed cell viability (Figure 2C) using CalceinAM (green) and propidium iodide (PI, red) as live/dead markers, respectively (Figures 2C and 2D). Astrocytes were highly viable in 3D-Fibrin (∼90%; 1.6 mM Ca^2+^, from AM), and showed a non-significant minor reduction in 3D-FAR at the concentration used for crosslinking (∼80%, 175 mM Ca^2+^;*p* = 0.0659; Figure 2E). To further evaluate the impact of Ca^2+^ on cell behaviour, we assessed astrocyte morphology in 3D-Fibrin hydrogels treated with the same Ca^2+^ concentration and time as in 3D-FAR: remarkably, we found a similar proportion of ramified cells in absence or presence of Ca^2+^ (3D-Fibrin ∼8%, 3D-Fibrin+Ca^2+^: ∼5%, *p* = 0.4392; Figures S1G-S1K). Furthermore, whole hydrogel imaging showed morphologically ramified astrocytes throughout the 3D-FAR hydrogel, suggesting a homogenous hydrogel microstructure and crosslinking throughout the hydrogel (Video S7). Together, our analyses indicated that 10 minutes of Ca^2+^ exposure required to crosslink alginate-RGD in 3D-FAR does not have major impact on astrocyte viability or ramification, implying that the spatial microstructure and higher stiffness of 3D-FAR hydrogels may be instrumental to shape astrocyte morphology in such 3D environment.

To analyse astrocyte proliferation on 2D and in 3D-Fibrin/3D-FAR hydrogels, cells were treated with the DNA-base analogue EdU twice, namely at the time of embedding and after 48h, and analysed 2 days later. Interestingly, astrocytes in 3D environments proliferated less than in 2D cultures, least in 3D-FAR (Figures 2G and 2H), resulting in a lower cell density in 3D-FAR already at 96 hours (Figure 2I, *p* = 0.0007). We also tested the impact of fibrin on astrocyte behaviour on 2D-coated glasses: Interestingly, astrocytes seeded on 2D-POL or on 2D-Fibrin showed similar morphology (Figures S1L and S1M), proliferation and density (Figures S1N-S1Q), thus suggesting that fibrin *per se* is not sufficient to explain the differences observed when cells are cultured in 3D-Fibrin, but rather the 3D environment substantially contributes to the proliferation behaviour.

Despite the lower proliferation and density, astrocytes embedded in 3D-FAR could be cultured in astrocyte medium (AM) containing leukemia inhibitory factor (LIF) over several weeks (Figures 2J-2L), maintained the expression of GFAP, and their processes reached other cells in dense areas after 14 days (Figure 2K). Conversely, at the conditions tested (2 million cells/ml), 3D-Fibrin hydrogels dissolved after 7 dpe. At higher density (5 million/ml), 3D-Fibrin hydrogels could be cultured for more than 3 weeks, mostly due to a dense cell layer forming on top of the hydrogels, sealing it from erosion (see section “Direct Neuronal Reprogramming in 3D”).

Taken together, our results suggest that a modest decrease in viability, combined with a pronounced reduction in proliferation, may account for the observed lower cell density in 3D-FAR compared to 3D-Fibrin hydrogels after 96h (Figure 2I).

### RNA-sequencing reveals reactive astrocyte state in FAR, and a cycling progenitor state in fibrin

#### 2D vs 3D

As astrocytes cultured on 2D-POL and 2D-Fibrin showed similar phenotype (Figures S1l-S1P), we decided to investigate the impact of the 3D environments at the molecular level by comparing the transcriptome of astrocytes cultured for 7 days either on 2D-Fibrin or in 3D-Fibrin, thus maintaining the same substrate and reducing possible confounding factors (Figure S2A). Interestingly, several hundred genes were differentially expressed (Figure S2B; log2fold change >1 and pval<0.05; Table S1), despite the heterogeneity of biological replicates (Figure S2C). Known astrocyte markers ^19^ were similarly expressed on 2D-Fibrin and in 3D-Fibrin (Figure S2D). As fibrin can interact with several components of the extracellular matrix, such as integrins ^36,37^ and collagen ^38^, we wondered whether a different environment (2D vs 3D) might affect their expression: indeed, some detected integrins and collagens were significantly more expressed in a specific condition (e.g., *ITGA2*, *ITGA3*, *COL7A1*, *COL4A5* and *COL4A6* in 2D, *ITGA9*, *ITGBL1*, *COL26A1*, *COL19A1* in 3D; Figures S2E and S2F, respectively), thus indicating a relevant impact of the environment. Gene ontology (GO) analysis on genes more expressed in astrocytes embedded in 3D-Fibrin (287 genes, pval<0.05; Figure S2B) revealed the presence of genes associated to generic terms as ‘*animal organ morphogenesis’* (Figure S2G, Biological Process, BP; Table S2), but also more specific, such as ‘*extracellular matrix’* (Figure S2H, Cellular Compartment, CC; Table S2) and ‘*peptidase regulator activity*’ (Figure S2I, Molecular Function, MF; Table S4). Conversely, genes related to ‘*inflammatory response’* (Figure S2J, BP; Table S5), ‘*cell periphery*’ (Figure S2K, CC; Table S6) and *’cytokine activity*’ (Figure S2L, MF; Table S7) were more expressed by astrocytes on 2D-Fibrin (Figures S2BC-G; 502 genes), suggesting a pronounced reactive state. In both conditions, the extracellular compartment was identified as significantly changed, but different genes were regulated (e.g., *MMP7*, *LGALS9*, *LAMB3*, *LAMA5* on 2D-Fibrin; *MMNR1*, *LAMC3*, *GPC3* in 3D-Fibrin; Table S2 and Table S6). Overall, transcriptome comparison revealed a substantial impact of the 3D over the 2D environment, even when astrocytes are cultured in or on chemically identical substrates.

#### 3D-FAR vs 3D-Fibrin

To evaluate the impact of different 3D substrates on human astrocytes, we assessed their transcriptome after seven days in 3D-Fibrin and 3D-FAR hydrogels (Figure 3A) and found extensive changes in gene expression (498 more expressed in FAR, 530 in fibrin; log2Fold change>1; pval<0.05; Figures 3B and Table S8), despite some variability across biological replicates (Figure 3C). GO analysis on genes more expressed in 3D-Fibrin revealed an enrichment for genes associated to proliferation (e.g., *MKI67*, *HMGB2*, *TOP2A*, *CENPE*, *CENPF*; Figures 3D for BP, S3A and S3B CC and MF; Tables S9-S11). Indeed, we detected a trend of more astrocytes (S100β^+^ cells) positive for HMGB2 ^39^ in 3D-Fibrin than in 3D-FAR (Figures S3C and S3D), both supporting the observation of their higher proliferation in 3D-Fibrin than in 3D-FAR (Figure 2F). Conversely, genes expressed at higher levels in 3D-FAR were associated to response to stimuli (e.g., *C3*, *CHI3L1, CHI3L2*, *CD44, LCN2, LGALS3, LGALS1, ATF3;* Figures 3B, 3C, 3E; Table S12), secretory vesicles and cytokine activity (Figures S3E, S3F; Tables S13, S14, respectively), thus suggesting a different reactive state, similar to 2D cultures. Indeed, on 2D-POL and in 3D-FAR we found a similar high proportion of S100β^+^ astrocytes positive for CHI3L1, a secreted protein impairing neurogenesis ^40^, and much higher than in 3D-Fibrin (Figures S3G, S3H), thus supporting a distinct reactive state of astrocytes in 3D-FAR than in 3D-Fibrin. As CHI3L1 is also part of human astrocyte maturity gene signatures ^41^, we compared astrocytes in 3D-FAR and 3D-Fibrin to genes associated to mature human astrocyte states ^41^ (Figure S3I): interestingly, CHI3L1 was the only gene significantly more expressed in 3D-FAR compared to 3D-Fibrin, thus indicating no overall difference in the maturation state between astrocytes cultured in the 2 different conditions. We then analysed the expression of genes associated to different reactive states, either induced *in vitro* or *in vivo*. Remarkably, we found that 3D-FAR astrocytes express high levels of genes induced by IL-1β treatment in human astrocytes (Figure 3F) ^42^, markers for pan-astrocyte reactive state ^43^, and genes ^43^more expressed in Alzheimer’s patients’ isolated astrocytes ^44^ (Figures S3J and S3K, respectively). Conversely, the transcriptome of 3D-Fibrin astrocytes was characterized by the presence of several genes identified in a model of middle cerebral artery occlusion (MCAO) in a cluster of astrocytes with moderate infarction (Figure 3G; cluster AC_3 ^45^), thus suggesting a different type of reactive state. Remarkably, few integrins and collagens were differentially expressed between 3D-Fibrin and 3D-FAR astrocytes (Figures S3L and S3M, respectively), suggesting an overall similar interaction capability of astrocytes in the different environments, which offered the same concentration of fibrin matrix (3 mg.ml^-1^). As ECM-associated proteins were overall not changed, we looked at the expression of metalloproteases (MMPs), which are capable of degrading a variety of extracellular matrix components^46^: interestingly, *MMP3*, *MMP7* and *MMP12*, which have been implicated in neuroinflammation ^47,48^, were more expressed in 3D- FAR, further supporting 3D-FAR as a pro-inflammatory environment (Figure S3N). Furthermore, given the different stiffness between 3D-Fibrin and 3D-FAR, we wondered whether mechanosensitive coding genes could be differentially expressed: indeed, two transient receptor potential genes, TRVP4 and TRPV2 were significantly differentially expressed (Figure S3O). Both are indeed expressed in astrocytes ^49,50^, with the former important to regulate cell volume ^51^, while the latter induced upon oxygen-glucose deprivation/reoxygenation (OGD/R) and reducing the expression of NGF, thus affecting neuronal survival ^52^.

**Figure 3.**
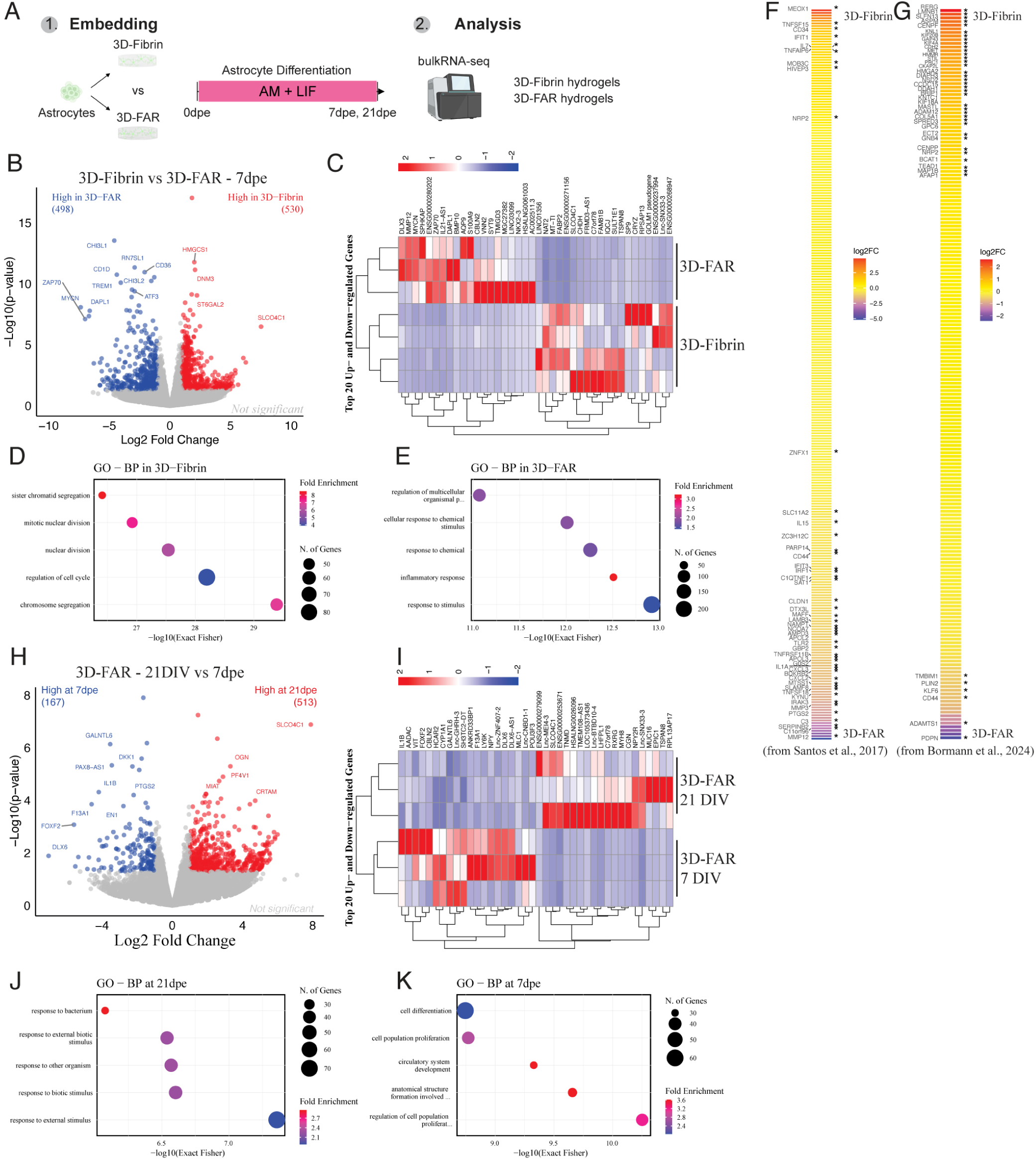
RNA-sequencing analysis of astrocytes embedded in 3D-Fibrin and 3D-FAR hydrogels. (**A**) Experimental design. Astrocytes were cultured in different gels and collected for RNA-seq at different time points. (**B**) Volcano plot showing differentially expressed genes (3D-Fibrin, red; log2FC>1, pval<0.05; 3D-FAR, blue; log2FC< -1, pval<0.05). Names are indicated when log2FC> abs(2). (**C**) Heatmap showing the top 40 genes more expressed by astrocytes in 3D-Fibrin or 3D-FAR. (**D, E**) Gene ontology (GO) analysis of biological processes (BP) from genes more expressed in 3D-Fibrin (log2FC>1, pval<0.05) or in 3D-FAR (log2FC < -1; pval < 0.05). (**F, G**) Heatmaps showing the log2FC of genes induced in human astrocytes upon IL-1β treatment (Santos et al., left panel) and more expressed in astrocytes following MCAO (Bormann et al., right panel). **(H)** Volcano plot showing differentially expressed genes in 3D-FAR at different time points (21dpe, red; log2FC>1, pval<0.05; 7dpe, blue; log2FC< -1, pval<0.05). Names are indicated when log2FC> abs(2). **(I)** Heatmap showing the top 40 genes more expressed by astrocytes in 3D-FAR at 7 dpe or 21 dpe. **(J, K)** GO analysis of BP terms from genes more expressed in 3D-FAR at 21 dpe (log2FC>1, pval<0.05) or at 7 dpe (log2FC < -1; pval < 0.05).

Overall, the comparison of 3D-Fibrin vs 3D-FAR astrocytes revealed significant differences in the proliferation capacity and reactive state, suggesting their use to investigate specific aspects of astrocyte biology and to use the 3D biomaterial environment to elicit different astrocyte phenotypes.

#### 3D-FAR short vs long term culture

To assess how long-term culture influences astrocytes in 3D-FAR and evaluate possible changes in the inflammatory state, we compared the transcriptome of cells cultured for 7 or 21 dpe (Figures 3H and 3I). Among the genes more expressed at 21 dpe (513; log2fold change>1; pval <0.05; Figures 3H and 3I; Table S15), several ones related to ‘*cell response*’ (Figure 3J; Table S16), ‘*plasma membrane*’ and ‘*ion channels*’ (in particular, K^+^ or Na^+^ channel-coding genes; Figures S3P, S3Q; Tables S17 and S18); conversely, genes more expressed at 7 dpe (167; log2fold change>1; pval <0.05; Figures 3H) were associated to ‘*cell proliferation*’ – including both pro- (e.g., *IL1A*, *IL1B*) and anti-proliferation genes (*BATF2*) – ‘*membrane microdomain*’ and ‘*extracellular space*’ (Figures 3K, S3R and S3S; Tables S19, S20, S21), thus indicating substantial changes of the astrocytes over time. Then, we wondered whether the inflammatory state found at 7 dpe persisted at 21 dpe: Strikingly, despite the observed biological variability, all the genes associated to the GO term ‘inflammatory response’ in 3D-FAR at 7 dpe (Table S12) were less expressed at 21 dpe (Figure S3T), while other genes associated to chronic inflammation ^12^ were expressed at similar level with the notable exception of genes associated to high inflammatory state, namely *IL1β*, *C3* and *CCL2* (Figure S3U).

Overall, the analysis of the transcriptome not only revealed that the 3D environment alters astrocyte behaviour compared to 2D, but also that 3D-Fibrin astrocytes are more proliferative, while 3D-FAR hydrogel maintains the cells in a reactive state, which is modulated over time.

### Distinct effects of 3D-FAR and 3D-Fibrin on direct astrocyte-to-neuron reprogramming

Our results suggested substantial differences between human astrocytes embedded in 3D-Fibrin and 3D-FAR. 3D-Fibrin is softer and fibrin rich, and the cells have a higher proliferation accompanied by a moderate reactive state, thus similar to intracerebral haemorrhage condition were astrocytes resume proliferation ^8,23^ and brain ECM changes ^15,53^ with an enrichment in fibrin; 3D-FAR is stiffer, astrocytes are less proliferative and showed a stronger reactive state. Therefore, we wondered if and how direct reprogramming of human astrocytes to neurons may be affected in these two very different 3D environments. To tackle this question, astrocytes were embedded in 3D-Fibrin or 3D-FAR together with retroviral vectors encoding either a fluorescent protein alone (DsRed) or coupled to a phospho-resistant ^54^ form of Neurogenin2 (9SA-Ngn2) ^55–57^, recently used to reprogram mouse ^58^ and human astrocytes ^19^ (Figures 4A and 4B). 2D-POL cultures, transduced with either control or 9SA-Ngn2 encoding retrovirus, were used as controls (Figure 4B). To mitigate the impact of cell death during the conversion process ^19,22^, we increased cell concentration to 5 mio.ml^-1^ in 3D (Figure 4A): this allowed to culture astrocytes not only in 3D-FAR (Figures 2F and 2G), but also in 3D-Fibrin for up to 21 days (Figure 4).

**Figure 4.**
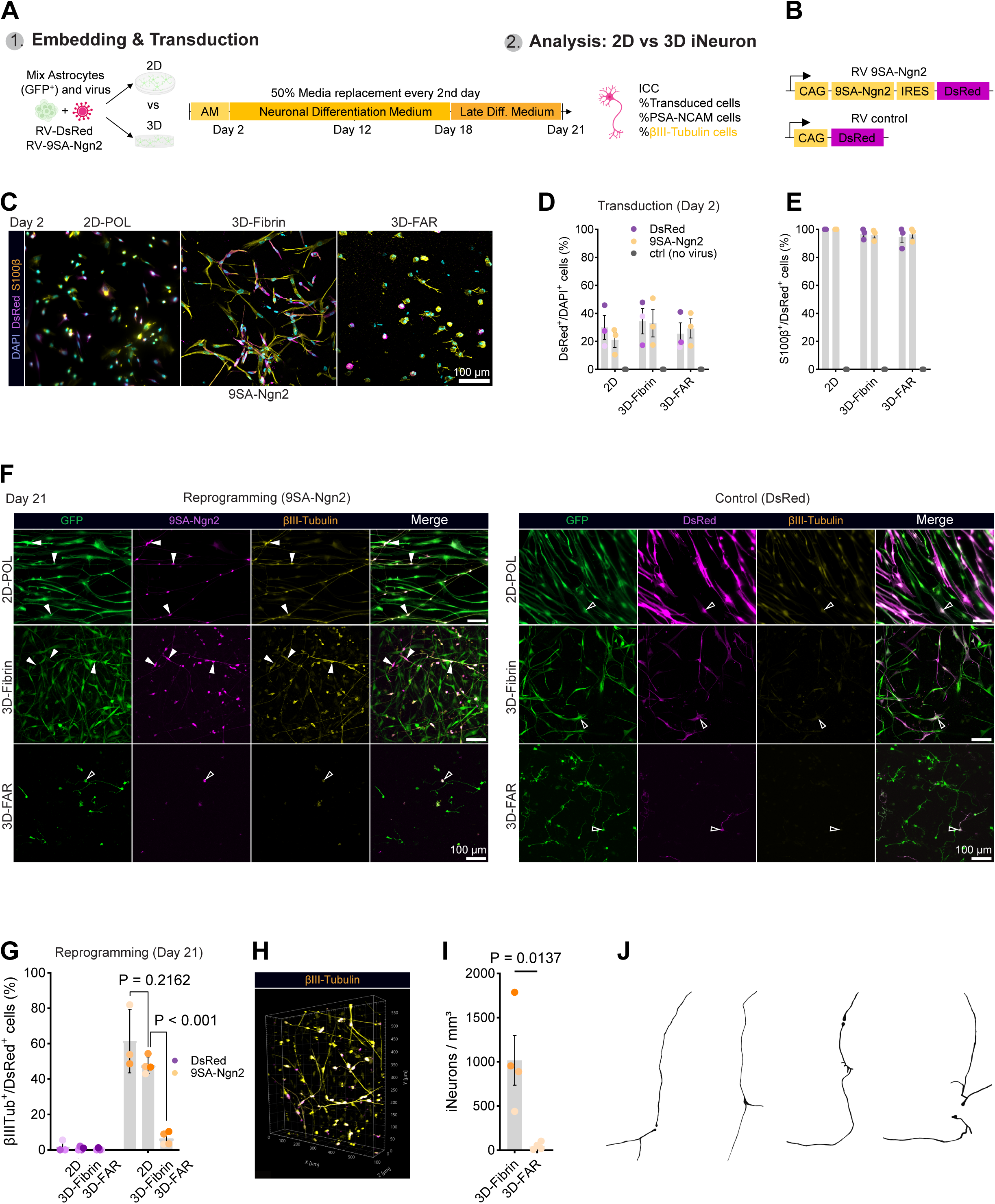
Direct neuronal reprogramming of astrocytes in 3D hydrogels (A, **B)** Experimental design for direct neuronal reprogramming in 3D (A) and scheme of the retroviral vectors used for direct reprogramming (B). (**C-E**) Confocal images of DsRed^+^ viral transduced cells after 2 dpi (C), transduction efficiency for both viruses in 2D and 3D cultures (D) and proportion of S100β astrocytes transduced (E) (N = 3(2), mean ± S.E.M). Scalebar = 100µm. (**F**) Fluorescence microscopy images (z-stack) of reprogramming performed either on 2D-POL or in 3D hydrogel cultures (Fibrin and 3D-FAR hydrogels). Scalebar = 100µm. (**G**) Graph showing the conversion rate 21 dpi (N = 3-4. Data shown as mean ± SD. Colors indicate quantifications form independent experiments. Un-paired student’s t-tests for individual comparisons). (**H**) Example 3D image of βIII-Tubulin^+^ cells in 3D-Fibrin hydrogel at 21 dpi. (**I**) Graph showing the density of induced neurons in Fibrin compared to 3D-FAR at 21 dpi (N = 4; data shown as mean ± SD. Colours indicate data from independent biological experiments. Un-paired student’s t-test). (**J**) Representative reconstructions of iNeurons in 3D-Fibrin (βIII-Tubulin^+^DsRed^+^); most iNeurons show simple uni- or bi-polar elongated neuron morphologies.

Two days after transduction (2 dpt), ∼33% of all cells were DsRed^+^, independent of 2D or 3D conditions (Figure 4C and 4D), and almost all were S100β^+^ (>97%, Figure 4E). After 21 dpt, very few astrocytes in 3D-FAR transduced with 9SA-Ngn2 had become βIII-tubulin positive and showed at least one thin primary process, indicative of a neuronal morphology (∼6%, Figures 4F and G). Conversely, both on 2D and in 3D-Fibrin 9SA-Ngn2-transduced astrocytes converted into *bona fide* neuronal cells (∼48-61% of DsRed^+^ cells were βIII-tubulin^+^ cells and had a neuronal morphology; Figures 4F and 4G), reaching a density of ∼1x10^3^ βIII-tubulin^+^ induced neuronal cells/mm³ in 3D-Fibrin (Figures 4H-J). This was significantly higher than when attempting reprogramming of astrocytes in 3D-FAR (*p = 0.0137*).

To check if 3D-FAR astrocytes could not be reprogrammed or may not survive until 21 dpt, (Figure 2K), we analysed the cultures undergoing direct conversion earlier at 12 dpt (Figure S4). Interestingly, we found that PSA-NCAM (neuronal marker expressed by newborn neurons) was expressed by a high proportion of control- and 9SA-Ngn2 transduced astrocytes (Figure S4A-C; 40-60% in controls; in 9SA-Ngn2 2D: ∼96%, 3D-Fibrin: ∼99%, 3D-FAR: ∼82%), but at significantly different level (Figure S4D; *p = 0.0100*). Remarkably, in 3D-FAR hydrogels a slightly lower proportion of PSA-NCAM^+^βIII-tubulin^+^/DsRed^+^ cells could be detected at 12 dpt (Figures 4A and E) and significantly lower compared to 2D culture (*p = 0.0184*). These transduced cells in 3D-FAR also appeared much smaller and had much shorter processes compared to 3D-Fibrin, thus suggesting that the astrocytes in the 3D-FAR substrate were impaired in their fate conversion, possibly leading to cell death (Figures 4C and 4F). Consistent with this, we observed many round cells with pyknotic DAPI^+^ nuclei among DsRed^+^ cells in 3D-FAR, explaining the lower cell density of induced neurons in 3D-FAR at 21 dpt. In sharp contrast, a high proportion of 9SA-Ngn2-transduced cells seemed already converted into neuronal cells at 12dpi in 3D-Fibrin, as evident by their morphology (Figure S4A) and by the expression of βIII-tubulin^+^ (Figure S4E).

Overall, human iPSC-derived astrocytes can be successfully reprogrammed into neuronal cells in 3D-Fibrin hydrogels, where we observed uni- or bipolar elongated morphologies (Figure 4J). Direct neuronal conversion was worse in stiffer 3D-FAR hydrogels, where astrocytes showed a stronger inflammatory signature after seven days (Figures 3C and 3K), highlighting the relevance of the 3D environment for direct neuronal reprogramming, and permissiveness of 3D-Fibrin environment to allow for conversion of human astrocytes into neurons.

## Discussion

### Distinct astrocyte phenotype in 3D-Fibrin versus FAR

We show here that the choice of 3D substrate can drastically influence human iPSC-derived astrocyte morphology, phenotype and reprogramming potential, as monitored by proliferation, migration, gene expression and direct conversion experiments. Astrocytes embedded in 3D-Fibrin retained a rather simple morphology (3-5 primary branches), and this is associated to a proliferative state, as found by live imaging, gene and protein expression analysis. Conversely, astrocytes in 3D-FAR showed a rather complex morphology with up to 20 branches – similar to astrocytes grown in glia-enriched 3D cortical organoids with 10-15 primary branches ^59^ – which was surprisingly accompanied by a different, more inflammatory reactive transcriptional signature. Human astrocytes extended their processes along the 3D-Fibrin structure (Video S2) and showed migratory behaviour and degradation of the hydrogels, as shown in other gel systems like fibrin ^16,17,60^, pristine collagen ^61^ or derivatives. Conversely, hydrogel degradation and migration were mitigated by using collagen crosslinked with 4S-StarPEG ^61^ and in our 3D-FAR gel. This allowed also for longer cultures and was associated with a more complex morphology of the astrocytes and higher expression of genes elicited by interleukin IL-1β treatment or present in Alzheimer’s disease. Further studies will identify which aspects of these substrates are responsible for the differences observed in astrocyte phenotypes. As astrocytes in pure alginate-RGD (Figure S1) did not show pronounced branching and a low degree of substitution with RGD peptides was provided on the alginate matrix in 3D-FAR (0.375% DOS; GRGDSP), it is unlikely that such small increase in RGD concentration lead to the observed changes; rather a synergistic effect of fibrin and alginate-RGD may act in shaping astrocyte behaviour, possibly connected to substantial changes in stiffness and porosity. Interestingly, continuous live-imaging in 3D-FAR showed that primary branches of astrocytes grow preferably along the AF594^+^ fibrin fibres (Video S2).

Noteworthy, the apparent elastic modulus K_eff_ measured via AFM in 3D-Fibrin hydrogels (∼300Pa) was in a similar range as reported in human non-malignant gliotic tissue ^32^ (∼10-180 Pa), while in 3D-FAR hydrogels K_eff_ was much higher (∼5 kPa) and more similar to the values measured in human glioblastoma multiforme ^32^(∼3kPa - 13.5 kPa), and indeed stiffer than physiological brain tissue (∼1000 Pa, fresh human tissue; or in bulk-tissue testing shear modulus G = 300-500 Pa, human brain tissue, *post mortem* time < 24h ^62^). It is important to note that stiffness does not suffice to describe the complex mechanical properties of brain tissue ^63^, and stiffness measurements differ profoundly depending on the technique used (e.g. AFM, magnet resonance elastography [MRE], rheology), across scales, loading modes (compression, tension, shear ^62^), speed, *post mortem* time, and tissue regions, such that a direct comparison to the *in vivo* situation is difficult. Yet overall, fibrin together with alginate-RGD peptides and much higher stiffness resulted in increased astrocyte branching at the expense of migration and proliferation.

The analysis of proliferation and transcriptome of astrocytes in 3D-Fibrin and 3D-FAR revealed remarkable differences (Figures 2 and 3): cells in 3D-Fibrin were more proliferative, similar to mouse astrocytes cultured on soft substrates (∼ 300 Pa)^64^ and showed a different reactive state compared to 3D-FAR. Indeed, a subset of astrocytes with this identity has been found in MCAO rats ^45^ and in lesions with intracerebral haemorrhage in mice ^20,65^ and men ^8,23^. Conversely, the reactive signature of the astrocytes cultured in 3D-FAR hydrogels showed similarities to tumour-associated astrocytes ^41,66^, human astrocytes treated with IL-1β ^42^, and with human Alzheimer’s astrocytes (Figures 3F, G, S3I, S3J), as those conditions showed a higher expression of several inflammation-associated genes (e.g. *CHI3L1, CHI3L2, SLC7A& S100A4) ^41,44,66,67^*. Remarkably, the inflammatory signature is present on 2D-POL and in 3D-FAR, both much stiffer than 3D-Fibrin, and it changes over time in 3D-FAR (Figures S3R and S3R), suggesting the adaptation of the astrocytes to the 3D environment.

Overall, 3D-FAR hydrogel shared several features with the peritumoral regions of human glioblastoma ^66^, such as elevated stiffness and pronounced reactive gliosis.

This supports the use of such hydrogels to model a strongly adverse microenvironment.

### Distinct astrocyte phenotypes differ profoundly in their capacity to convert into neurons

The functional relevance of these distinct astrocyte phenotypes elicited by the different 3D environments was demonstrated by their huge difference in neuronal reprogramming: interestingly, 3D-FAR embedded astrocytes, transduced with 9SA-Ngn2, expressed the neuronal markers PSA-NCAM and βIII-tubulin at 12 dpt. However, already at that time many cells were round, indicative of an unhealthy state, and virtually no converted neurons could be detected under these conditions at 21 dpt. Thus, most of the cells undergoing direct conversion might have reverted to an astrocyte identity or succumbed to cell death, a more likely possibility given the striking difference in cell density observed at 21 dpt (Figure 4F). Our 3D-FAR model is thus ideally suited to explore means to overcome the reprogramming block of this astrocyte state with an inflammatory signature, and revert their phenotype to a healthier state. Conversely, astrocytes in 3D-Fibrin readily reprogrammed into neuronal cells when transduced with the 9SA-Ngn2, with similar proportions to the classical 2D substrates described before^19^. While the direct conversion of human fibroblasts into induced neurons in 3D has already been shown ^68–70^ and recently demonstrated for human glia spheroids ^71^, we revealed for the first time how the specific ECM environment alters cell state and significantly contributes to shape the direct conversion of human astrocytes into neuronal cells, which occurs in soft fibrin but not in stiffer 3D-FAR substrate.

An *in vivo* analogous astrocyte state exists with the astrocytes resuming proliferation after injury and partially dedifferentiating to an earlier progenitor fate ^11,23,72^. Indeed, targeting viral vectors to this population allows efficient reprogramming in murine models of traumatic brain injury ^22^. However, neuronal cells resulting from direct reprogramming in 3D-Fibrin showed simple morphologies with hardly any branching and a single long primary process (Figure 4G). This, indeed, represents a novel model for future studies aimed at investigating the differentiation, maturation, and connectivity formation of newly generated neurons in 3D, and identify conditions (e.g., substrates, co-cultures or co-factors) leading to the generation of fully functional 3D-embedded neuronal circuitry – likely being facilitated by increasing to higher cell density ^71^, survival of converting cells, co-cultures, and higher viral titre.

Taken together, our work not only shows how different fibrin-containing substrates can elicit very distinct astrocyte states, revealing how this affects direct conversion into neurons, but also provides new model systems compatible with high content screening for overcoming the inflammatory reactive astrocyte state resistant to reprogramming and optimizing reprogramming outcome. As 3D-FAR showed stiffness in the range of human grade IV glioblastoma^32^, with astrocytes expressing gene signatures related to human tumor-associated astrocytes^66^, and simultaneously elicited mitigated reprogramming, it may represent an adequate 3D *in vitro* model to study further the reprogramming of glioblastoma-associated astrocytes into neurons in the context of cancer reprogramming in stiff tumor environments ^73^. Moreover, co-culture with neurons or other cell types may allow to determine the effects of the astrocyte state in 3D-FAR conditions on neuronal function and synaptic communication upon direct conversion ^68^.

### Limitations of the study

A fascinating topic in astrocytes biology is their heterogeneity across and within CNS regions: the protocol employed to differentiate iPSC into astrocytes does not contain a pre-patterning step, and consequently, the resulting astrocytes do not show any specific regional identity. Future studies will address whether patterned astrocytes behave differently in similar 3D environments. Likewise, we did observe various morphologies of astrocytes, mostly in the 3D-FAR condition. This raises the questions if such heterogeneity depends on the cellular density or on the biomaterial environment properties (stiffness, microstructure), implying a relevant role of local contacts or microenvironment differences. Hence, this could offer a potential reference to the astrocyte heterogeneity developing *in vivo*, or whether it already emerges during the differentiation of iPSCs into astrocytes or by providing different 3D biomaterial-properties to the cells alone. In future work we will disentangle microstructure (e.g. via electron microscopy) from stiffness to explore further the role of single biomaterial properties on cell heterogeneity. Furthermore, future studies will investigate the direct reprogramming of astrocytes within distinct 3D hydrogel environments using single-cell analyses to elucidate their contribution to shape the molecular mechanisms of direct neuronal reprogramming.

## Acknowledgements

We thank Dr. Andreas Thomae and Dr. Mariano Pisfil from the BMC Bioimaging facility for supporting two-photon microscopy and live-cell imaging, Dr. Madeleine Schmitt und Dr. Jörg Renkawitz for plasma coater access. Library preparation and sequencing was performed at the Helmholtz Zentrum München (HMGU) by the Genomics Core Facility (Dr. Gertrud Eckstein, Dr. Inti De la Rosa Velazquez). We acknowledge the technical support of Core Facility Genomics at Helmholtz Munich. We thank Dr. Thomas Walzhoni for help with the Galaxy workflow pipeline for RNA-seq data alignment and processing. We thank Dr. Julia Becker (Cambridge, PDN) for Sneddon model fitting in AFM experiments. We thank Dr. Maria Richter for suggestions on bioinformatic analysis. We thank Claire Malvezin for support in calcium exposure and 2D comparison assays. We would like to thank PD Dr. Swetlana Sirko for discussions on astrocyte biology and Dr. Yvette Zarb for critical discussion on the manuscript. This work was funded by the German Research Foundation TRR274 (no. 408885537, M. Götz), FOR2879/2 (no. 405358801, M. Götz), and SyNergy (EXC2145/Project-ID 390857198, M. Götz). Parts of this work were supported by a New Frontiers in Research Fund Transformation grant to M. Götz, funded through three Canadian federal funding agencies (CIHR, NSERC, and SSHRC), the European Union’s Horizon 2020 research and innovation program under grant agreement no. 874758 (NSC Reconstruct to M. Götz).

## Author contributions

Conceptualization: G.M., T.D., and M.G.; methodology: T.D. (hydrogels, 3D cell culture, live-imaging), T.D., G.M., M.G. (3D reprogramming), M.B. and G.M. (iPSC-to-astrocytes differentiation), M.B. (virus production), SVS and K.F. (AFM); investigation: T.D. and M.B. (reprogramming experiments), T.D. and S.V.S. (AFM experiments), T.D. and K.K.D. (bulk RNA-seq experiments, proliferation and viability assays); resources: K.F. and M.G.; formal analysis: T.D., K.K.D., S.V.S., M.R., G.M.; supervision: G.M. and M.G; funding acquisition: K.F. and M.G.; writing: T.D., G.M., M.G., and all authors contributed corrections and comments.

## Methods

### Human iPSC culture

iPSCs were cultured similarly to described before ^19^. Briefly, human iPSCs were expanded on 6-well plates (Sarstedt) coated with Geltrex^TM^ (LDEV-Free Reduced Growth Factor Basement Membrane Matrix, Thermo Fisher, A1413302) in mTESR1 medium containing mTESR1 supplement (hiPSC medium). The medium was changed daily; passaging was performed via collagenase Type IV (Stem Cell Technologies, 07909) treatment (37°C, 5-7 minutes). Collagenase was aspirated and fresh medium was added to each well. Cells were collected using a cell scraper, followed by plating the cells on a new 6-well plate at a desired density (Split ratios: 1:4 or 1:8). We used iPSC or generated transgenic iPSC constitutively expressing green fluorescent protein (GFP), generated by Piggybac integration and selection.

### Proliferating astrocyte differentiation from human iPSC

Human proliferating astrocytes (“astrocytes”) were generated as previously described ^19^. In brief, human iPSC were dissociated (collagenase IV) and cultured in suspension culture to form embryoid bodies. Next, cells were cultured for 24h in hiPSC medium supplemented with Rock Inhibitor (10 mM, Y-27632, Stem Cell Technologies, 72304). On the next day, medium was changed to astrocyte medium (AM, ScienCell, 1801) containing supplements (10 ng.ml^-1^ PDGF-AA, R&D Systems, 221-AA; 20 ng.ml^-1^ Noggin, Peprotech, 120-10C) in which the cells were cultured for 14 days, followed by another 7 days in AM containing just PDGFAA. On day 21, the embryoid bodies were dissociated with a pipette and astrocytes were seeded on poly-L-ornithine (PLO) and laminin (LAM) coated petri dishes. Astrocytes were cultured in AM containing endothelial growth factor (10 ng.ml^-1^, recombinant human EGF, Peprotech, AF-100-15) and fibroblast growth factor (10 ng.ml^-1^, recombinant human FGF-basic, Peprotech, AF-100-18B) until day 35 (d35), with media changes every second day, and passaged using Accutase (StemPro^TM^ Accutase®, A11105-01) when reaching 80% confluency.

### 3D cell culture of human astrocytes in fibrin hydrogels

Fibrin hydrogels (final concentration: 3 mg.ml^-1^) were prepared similarly to as described before ^74^ by combining equal amounts (v/v) of a solution A, containing fibrinogen (6 mg.ml^-1^, from bovine plasma, Sigma Aldrich; 341573) and aprotinin (80 U.ml^-1^, Roche, 10236632103) in 1X PBS, with a solution B containing thrombin in AM (4 U.ml^-1^, Sigma Aldrich; T4648). Stock aliquots of fibrinogen, thrombin, and aprotinin were stored at -21°C and freshly thawn on the day of the experiment. Fibrinogen stock aliquots were thawn for 30 minutes at 37°C. Before hydrogel formation, aliquots were kept on ice prior to mixing solution A and B. For culturing astrocytes in 3D-Fibrin hydrogels, astrocytes (d35) were detached using accutase (3 ml, 7 min, per 10 cm petri dish) and counted using the direct trypan blue exclusion method using a Neubauer chamber (Carl Roth, T729.1). 400.000 cells were pelleted (300 rcf, 5 min) and redispersed in 100 µl of Solution B (4 mio.ml^-1^). Next, 50 µl of solution B were quickly mixed with 50 µl solution A, avoiding air bubbles, to ensure a homogenous distribution of the cells. 5 µl of cell-containing hydrogel precursor were casted into d = 5 mm mini-wells, which were custom prepared silicone wells covalently attached to glass coverslips (d = 12 mm) by plasma treatment ^75^. Gels were allowed to polymerise at 22°C (room temperature, RT) for 10 minutes. The final gels contained 3 mg.ml^-1^ fibrinogen, 40 µg.ml^-1^ aprotinin, 2 U.ml^-1^ thrombin, and 2 mio.ml^-1^ astrocytes. After polymerisation, gels were covered with 1 ml of AM and transferred into an incubator for subsequent cell culture.

### Culture in 3D alginate, Cultrex^TM^, and 3D-FAR hydrogels

#### Alginate

To culture astrocytes in 3D alginate-RGD hydrogels, low molecular weight sodium alginate pre-functionalized with RGD peptide by the manufacturer (Novatach, Cat# 4270101 VLVG, MW = 32.000g.mol^-1^, guluronic acid content > 60%, 0.376% degree of substitution [DOS], GRGDSP peptide) was used. Sodium alginate was dissolved in sterile 1X PBS to prepare a 2%(w/v) stock solution. Next, cellular pellet (200.000 cells) was redispersed in alginate precursor (0.5%(w/v)) and 12.5 µl were casted into custom made silicone mini-wells place in 24-well plates (2 mio.ml^-1^). To initiate crosslinking, hydrogel precursor in the mini-well was covered using a dialysis membrane (washed in ddH_2_O, cut into 9x9 mm squares, prior soaked in 175mM CaCl_2_ (prepared from Cat# 223506) solution). Hydrogels were covered with 1 ml of Ca^2+^ solution for 10 minutes to facilitate crosslinking of the alginate network. The initial covering using a dialysis membrane acted as a barrier to avoid unwanted initial mixing of the CaCl_2_ solution with the low viscosity alginate precursor in the mini-well. After 10 minutes, Ca^2+^ solution was removed, gels washed once with 1X PBS for 1 minute, and the cell-containing hydrogels were incubated in 1 ml of AM under standard incubation conditions (37°C, 5.0% CO_2_).

#### Cultrex^TM^

Cultrex basement membrane aliquots (Cultrex^TM^ 3-D BME, reduced growth factor, R&D Systems, 3445-005-01) were thawn on ice and pipetted for homogenization using cooled (4°C) pipette tips. Cell pellets were dispersed using the hydrogel precursor and pipetted (15 µl) inside silicone mini-well, followed by homogenous distribution via circular motion to prepare hydrogel discs. Cell-containing hydrogels were incubated for 30 minutes at 37°C in incubator to facilitate hydrogel crosslinking.

#### 3D-FAR

Fibrin-Alginate-RGD (3D-FAR) hydrogels were prepared similarly to fibrin hydrogels and without oxidation ^76^, using a low molecular weight alginate (Novatach VLVG alginate stock; 2%(w/v), MW: 40.000 Da). Alginate-RGD was diluted in solution A components to achieve final concentrations of fibrinogen (6 mg.ml^-1^), aprotinin (80 U.ml^-1^) and 1%(w/v) alginate, in 1X PBS. A typical hydrogel solution contained 50 µl alginate (2% w/v), 36.4 µl 1X PBS, 1.6 µl aprotinin, and 12 µl fibrinogen (Solution A: V_total_ = 100 µl, 1%(w/v) alginate, 6 mg.ml^-1^ fibrinogen, 80 U.ml^-1^ aprotinin). Solution A was homogenised using a pipette. Hydrogel formation was triggered by mixing equal amounts of solution A with solution B (4 U.ml^-1^ thrombin, in AM) containing the cells. The mixture was rapidly mixed by pipetting eight times, and the hydrogel mix (15 µl) was placed into the mini-wells. Final 3D-FAR gels contained the same amounts of fibrinogen, thrombin, and aprotinin as pure fibrin hydrogels, but additionally 0.5% (w/v) sodium alginate-RGD. Gels were allowed to gel for 10 minutes at RT. Then, hydrogels were covered with 175mM CaCl_2_ solution for another 10 minutes to crosslink the sodium alginate network inside the fibrin matrix, which resulted in interpenetrating network hydrogels (IPN). Gels were washed for 1 min with 1X PBS and transferred into respective media (AM+LIF; or reprogramming medium).

#### 2D-POL-Glass and 2D-Fibrin cultures

To compare the effect of different 2D substrates on astrocyte morphology and proliferation, astrocytes (d35) were seeded on glass coverslips coated with PO (20 µg.ml^-1^, in ddH_2_O, 16h overnight at 37°C; Sigma Aldrich; P3655) and LAM (10 µg.ml^-1^ in 1X PBS, min. 16h overnight at 37°C; Sigma Aldrich; L2020) to get 2D-POL-coated glass ^19^, and compared to astrocytes seeded on 2D-Fibrin hydrogels. Briefly, 2D-Fibrin hydrogels were casted into mini-wells by adding 12.5 µl of fibrin hydrogel (3 mg.ml^-1^) distributed evenly in the wells and allowed to polymerise for 10 min at RT to create flat hydrogel discs. The hydrogels were prepared the same way as 3D-Fibrin, but without embedding astrocytes (see “3D cell culture of human astrocytes in fibrin hydrogels”). Finally, astrocytes (35.000 in 1 ml AM+LIF, per 24-well) were seeded either on top of 2D-POL-Glass or on 2D-Fibrin hydrogels, which were prior placed into 24-well plates, and cultured for 96h or 7 days.

### Mechanical testing by atomic force microscopy (AFM)

To assess the mechanical properties of Cultrex^TM^, 3D-Fibrin, and 3D-FAR hydrogels, we performed AFM at 37°C immediately after hydrogel fabrication (∼20 minutes for each gel; N = 3 for all conditions) from two independent batches of fibrinogen (Sigma Aldrich; 341573, #Lot1 3897584; #Lot2 4177330) in case of 3D-Fibrin and 3D-FAR. In detail, for Cultrex^TM^ three gels from individual preparations were measured. For 3D-Fibrin and 3D-FAR hydrogels, a total of six gels were cast (three individual preparations for each #Lot) for each type of gel. AFM measurements were conducted on a JPK CellHesion 300 atomic force microscope (JPK Instruments), mounted to an upright setup with an Axio Zoom.V16 (Zeiss) microscope with a pco.edge 4.2 bi sCMOS camera. Tipless silicon cantilevers (Arrow-TL1, NanoWorld) with a custom-attached polystyrene bead as probe (∼37 µm diameter, microParticles GmbH) with spring constant k (Arrow-TL1, NanoWorld) of 0.02 to 0.04 N/m, determined by the thermal noise method^77^, were selected. All samples were heated to 37°C using a PetriDishHeater (JPK Instruments) during all measurements. Brightfield images of the hydrogel surface before measurements were collected by a pco.edge 4.2 bi sCMOS camera connected to a Carl Zeiss Axio Zoom.V16 (Zeiss) microscope. Force-distance curves were recorded and analysed with the JPKSPM Data Processing software (version 8.0.192). First, a baseline subtraction with correction for offset and tilt was conducted on the last 20%, followed by a correction for cantilever bending. Finally, the Sneddon model for a spherical indenter (R = 18.64 µm) was fit. In order to not make any assumptions about the Poisson’s ratio ***ν***, ***ν*** was set to 0, yielding the reduced apparent elastic modulus K instead of the Young’s elastic modulus E. Stiffness values are reported as K.

### Microstructure analysis of fibrin and FAR hydrogels

The porosity and microstructure of 3D-Fibrin and 3D-FAR hydrogels were imaged using a Leica Stellaris 8 FALCON system (20x objective). 3D-Fibrin and 3D-FAR hydrogel fibres were visualised by substituting 1 µl of fibrinogen with 1 µl of Alexa Fluor 594-conjugated fibrinogen (Thermo Fisher, F13193) into Solution A during hydrogel formation. The hydrogels were transferred to the microscope and imaged (field of view: 415x415x102 µm³, Voxel size: 0.4058x0.4058x2.0014 µm³). The microstructure was analysed from N = 3 independent hydrogel preparations. AF594^+^ hydrogel fibres were segmented using threshold adjustment in ImageJ, followed by quantifying the area between AF594^+^ hydrogel fibres.

### Immunocytochemistry

Astrocytes cultured in 3D hydrogels were washed once using 1X PBS and fixed with 4% PFA (in 1X HBSS) for 10 minutes at RT. Samples were washed three times (10 min each) with 1X HBSS followed by blocking and permeabilization for 1 hour at RT using 3% BSA, 0.5% Triton X-100 in 1X HBSS. Next, 3D structures were immunostained with the following primary antibodies: Mouse m1 anti-S100beta (Sigma, S2532, 1:1000), rabbit anti-glial fibrillary acidic protein GFAP (Sigma, G9269, 1:300), rabbit anti-YKL-40/CHI3L1 (Abcam, ab77528, 1:300), rabbit anti-HMGB2 (Abcam, ab67282, 1:1000), chicken anti-GFP (Aves Lab, 1020, 1:500), rabbit anti-RFP (Rockland, 600-401-379, 1:500), guinea pig anti-b-III-Tubulin (Synaptic systems, 302-304, 1:200), mouse anti-Polysialic Acid-NCAM (Merck Millipore, MAB5324, 1:300), in blocking solution, overnight at 4°C. The next day, samples were washed three times with 1X HBSS (15 min each) to remove the primary antibody solution, and then incubated with secondary antibody (1:500) (for details, see STAR methods key resource table) for 1.5 hours at RT in the dark. Samples were washed three times with 1X HBSS (10 min each), and DAPI (1:1000) was added to the second washing step to counterstain for nuclei when applicable.

### Morphology analysis

The morphology of astrocytes in 3D-Fibrin and 3D-FAR hydrogels was analysed using segmentation, reconstruction, and Sholl analysis, similar to described elsewhere ^59^. Astrocytes were cultured for 7 days in 3D-Fibrin or 3D-FAR hydrogels (2 mio.ml^-1^) and stained for S100β to identify astrocytes. Astrocytes embedded in the hydrogels were imaged using a Leica SP8 Falcon system in two-photon excitation mode (lasers: Insight X3 DUAL, 1045, Spectra Physics, with tuneable emission (680 - 1300nm). Pulse width <120 fs. Second laser: 1045 nm, pulse width <200 fs), equipped with a HC IRAPO L 25x/1.00 W motCORR (WD 2.6 mm) objective, with z-sectioning of 2 µm, 1024x1024 pixels, scan speed 400 Hz, dwell time >1.5 µs. Single astrocytes were selected from the imaging z-stacks using Fiji^78^ (ImageJ v1.54m), and astrocyte morphology was segmented and reconstructed by manual process tracing using neuTube ^79^ (v1.0z) to generate SWC files. Finally, astrocyte morphology was analysed from SWC files by Sholl analysis (radius step size: 2 µm) using the simple neurite tracer ^80^ (SNT) plugin (v4.2.1). Parameters like total neurite length, number of branches, and number of primary branches were considered for analysis of astrocyte morphology (n = 10 cells per condition, N = 3 biological experiments, n = 30 cells in total for each condition, 3D-Fibrin and 3D-FAR).

### EdU proliferation study

Edu (10 mM) was added to AM at day 0 of cell seeding and embedding in the hydrogels, and a second EdU treatment was added after 48h of incubation. Two days later, astrocytes were fixed and EdU detection was performed using the ClickiT^TM^ plus EdU cell-proliferation kit (Thermo Fisher, C10640), which is based on a copper-catalyzed azide–alkyne cycloaddition (CuAAC) of an azide-fluorophore with the alkyne containing EdU. EdU was visualised by an azide-Alexa Fluor 647 probe supplied in the kit. All steps were performed according to the manufacturer’s protocol. Nuclei were counter stained by DAPI (1:1000) incubation for 15 minutes at the end of the ClickiT^TM^ protocol, before the final washing with 1X HBSS. N = 4 biological experiments were performed. Astrocytes on 2D glass coated with PO+L served as control. EdU^+^ cells were quantified from three overview images (598.05x598.05x50 µm³) per condition per biological replicate. Cell quantification was performed via Fiji using the CellCounter (v3.0.0) plugin.

### Viability analysis

To test the influence of different Ca^2+^ concentrations on astrocytes viability, we performed a LIVE/DEAD cell viability analysis. In brief, we used CalceinAM (4 µl.ml^-1^, Thermo Fisher, C1430) and Propidium Iodide (1µl.ml^-1^, Sigma Aldrich, P4864) in 1X HBSS to indicate LIVE cells (CalceinAM conversion to fluorescent calcein upon cell metabolism) and DEAD cells (PI incorporation by cell permeability), respectively. Cells were cultured in 3D-FAR hydrogels crosslinked with either 50, 100, or 175mM CaCl_2_. astrocytes cultured in 3D-Fibrin hydrogels (1.6 mM CaCl_2_, from AM) served as control. After 96h of astrocytes cultured in AM+LIF, the cells were incubated for 45 minutes at 37°C, 5% CO2 in CalceinAM/PI solution, and directly transferred to a confocal microscope (Carls Zeiss LSM710 confocal) for imaging. Image stacks were recorded (N = 4 biological experiments, two technical replicates per condition, 485x485x50 µm³). Alive cells (Calcein^+^) and dead cells (PI^+^) were quantified using Fiji.

### Continuous live cell imaging

To investigate the direct interaction of cells with the different 3D environments (3D-Fibrin, 3D-FAR), we performed continuous live cell imaging of CAG::eGFP hiPSC-derived astrocytes for the first 72h of culture after embedding in the hydrogels. To visualise the hydrogel environment, we added 1µl of AF594-fibrinogen during hydrogel formation. LIVE-cell imaging was performed in a humidified chamber (TokaiHit STXG) at 37°C, 5% CO_2_, using a Leica Stellaris 8 FALCON system equipped with a 20X objective. Images were recorded every 30 minutes (field of view: 581x581x50 µm³) to avoid phototoxicity. Images were processed for further analysis using Fiji (ImageJ). 3D animation and reconstruction of confocal and multiphoton images was performed using Imaris (Oxford Instruments, x64, v9.6.0).

### Bulk RNA-sequencing

Astrocytes were cultured for 7 days (2D-Fibrin, 3D-Fibrin) and for 7 days and 21 days (3D-FAR) in the hydrogels and subjected to bulk RNA-seq analysis. RNA was isolated using the Arcturus PicoPure^TM^ RNA isolation kit (Thermo Fisher, KIT0204) following manufacturer instructions. Hydrogel discs containing the cells were scooped out from the mini-wells, homogenised in 100 µl of RNA extraction buffer in 500 µl Eppendorf tubes, briefly centrifuged and incubated for 30 minutes at 42°C. RNA samples were stored at -80°C until the subsequent RNA purification step. After purification, RNA quality was determined using a 2100 BioAnalyzer (Agilent) and the RNA 6000 pico kit (Agilent Technologies, 5067-1513). RNAs with a sample RIN value ≥ 7.9 (mean RIN = 8.7 ± 0.6) were selected for mRNA sequencing (poly-A selected). RNA quantity was determined using Qubit™ RNA High Sensitivity (HS) kit, and 10 ng of RNA (in 10.5 µl non-DEPC treated H2O) were pipetted into a Greiner bio sapphire microplate 96-well (Greiner Bio-one, 652290). Library preparation was performed using the SMART-Seq mRNA LP kit (Takara, 634769) starting with 10 ng total RNA, following the kit’s instructions. After a final QC, the libraries were sequenced in a paired-end mode (2x100 bases) in the NovaseqX+ sequencer (Illumina) with a depth of ≥30 million reads per sample. RNA-seq raw data processing was performed in Galaxy (see workflow at Code Availability). Reads were trimmed using Trimmomatic ^81^ (v0.4) and the transcriptome was aligned against the human genome hg38 (GRCh38) reference, version 32 (Ensembl 98)^82^, using STAR aligner (v2.7.11b). We analysed three biological replicates (3D-FAR at 7 dpe and 21 dpe) and four biological replicates (2D-Fibrin and 3D-Fibrin) per condition.

### Bioinformatic analysis of bulkRNA-seq data

Data analysis or bulk RNA-seq data was performed in R Studio (v2024.04.2). Count matrices were TPM normalised. DEG analysis was conducted using DESeq2 (v1.44.0) analysis of the raw count matrix. Genes were considered differentially expressed with log2FoldChange > 1 (upregulated), or < -1 (downregulated), and pvalue < 0.05. Gene Ontology analysis was performed using TopGO ^83^ (Alexa A, Rahnenfuhrer J (2025). topGO: Enrichment Analysis for Gene Ontology. doi:10.18129/B9.bioc.topGO, R package version 2.59.0, https://bioconductor.org/packages/topGO). Heatmaps of DEG data and comparison to different astrocyte scores ^20,41,44^ were plotted using the pheatmap package (v1.0.12). Volcano plots were generated using the ggplot2.

### Direct neuronal reprogramming in 3D

To investigate the conversion of astrocytes into induced neurons (iNeurons), we reprogrammed astrocytes inside 3D-Fibrin and 3D-FAR hydrogels by mixing cells, viral particles, and hydrogel in one step: astrocytes (10 mio.ml^-1^) were redispersed in solution B (AM + thrombin) and mixed 50:50 with solution A containing fibrinogen, aprotinin, and virus (titre: 2.5x10^4-5^ viral particles/35000 cells in all conditions). In case of 3D-FAR, solution A contained sodium alginate-RGD as well (all in PBS; final cell concentration: 5 mio.ml^-1^). The volume of PBS was adjusted so that the addition of virus did not alter the concentration of the final hydrogel components (final concentration: 3 mg.ml^-1^ fibrinogen). We used a retrovirus CAG-9SA-Neurog2-IRES-DsRedExpress2 ^19^ (9SA-Ngn2) to express the phospho-resistant form of Neurogenin 2, or retrovirus CAG-DsRedExpress2 ^84^ as a control virus. Retrovirus vectors were produced ^19^, tittered ^85^ and used at a stock titre of 10^7^-10^8^ ml^-1^.The reprogramming paradigm was similarly performed as described before ^19^: astrocytes in 3D hydrogels and on 2D POL glass controls were cultured in neuronal differentiation media (DMEM/F12 + Neurobasal media [1:1], supplemented with 1x PS, 1x B27, 1x N2, and 1x MEM non-essential amino acids [NEAA]) which was supplemented with factors GDNF (2 ng/ml), CHIR99021 (2 mM), LDN-193189 (0.5 mM), LM-22A4 (2 mM), NT3 (10 ng/ml), SB-431542 (10 mM), db-cAMP (0.1 mg/ml), Noggin (50 ng/ml), and valproic acid sodium salt (VPA; 1 mM) ^19,86^. Every second day, half of the medium was replaced with fresh medium. Cultures undergoing direct reprogramming were maintained at 37°C, 5% O_2_, 5% CO_2_, in an incubator. On day 18, neuronal differentiation media was supplemented with db-cAMP, GDNF, LM-22A5, and NT3^19^. Cells were fixed on day 21 (4% PFA, 10 minutes), washed (three times 1X HBSS, 10 minutes), and stored at 4°C until further use. Transduced cells were considered successfully converted to induced neuronal cells when βIII-Tubulin^+^DsRed^+^ cells showed at least one thin process indicative of a neuronal morphology. To assess early conversion, cells were cultured for 12 days and early neuronal fate of transduced cells was quantified as PSA-NCAM^+^βIII-Tubulin^+^DsRed^+^ and PSA-NCAM^+^/DsRed^+^ cells. PSA-NCAM intensity of DsRed^+^ cells was quantified by measuring the mean PSA-NCAM fluorescence intensity of single cells per condition (3D-FAR: 93; 3D-Fibrin: 128; 2D: 103) using Fiji (ImageJ). Regions of interest (ROI) on maximum intensity projections of single cells were manually drawn, or by thresholding when applicable, followed by subtraction of the average image background intensity from 3x100µm² selections from individual images.

### Statistical analysis

All experiments were performed with at least N = 3 biological replicates per condition. Image analysis was performed blinded using the ‘Blind Analysis Tools’ file name encrypter toolbox (ZMBH Imaging Facility, University of Heidelberg) in ImageJ. All statistical tests are indicated at the presented figure captions. Statistical analysis was performed either using GraphPad Prism software (v10.4.1) or R (bulkRNA-seq data analysis; DESeq2). Pairwise comparisons of two conditions were performed using paired students t-test. Non-normally distributed data was analysed using Welch’s t-test. Comparisons between more than two conditions was performed using one or two-way ANOVA. For morphological analysis, statistical testing was performed on average values from the single cells measured in individual experiments (N = 3), and not on single cell events.

### Code availability

Galaxy workflows for raw RNA-seq data processing (paired-end sequencing, MultiQC report) are available upon request and will be available at the following Github repository (https://github.com/PhysiologicalGenomicBMC). Original R code used to process count matrices from RNA-seq data analysis is available upon request.

## References

1. Soto, J.S., and Khakh, B.S. (2023). Cell morphology: Astrocyte structure at the nanoscale. Preprint at Cell Press, 10.1016/j.cub.2023.01.045 10.1016/j.cub.2023.01.045.

2. Miller, S.J. (2018). Astrocyte heterogeneity in the adult central nervous system. Preprint at Frontiers Media S.A., 10.3389/fncel.2018.00401 10.3389/fncel.2018.00401.

3. Escartin, C., Galea, E., Lakatos, A., O’Callaghan, J.P., Petzold, G.C., Serrano-Pozo, A., Steinhäuser, C., Volterra, A., Carmignoto, G., Agarwal, A., et al. (2021). Reactive astrocyte nomenclature, definitions, and future directions. Preprint at Nature Research, 10.1038/s41593-020-00783-4 10.1038/s41593-020-00783-4.

4. Patani, R., Hardingham, G.E., and Liddelow, S.A. (2023). Functional roles of reactive astrocytes in neuroinflammation and neurodegeneration. Nat Rev Neurol 19, 395–409. 10.1038/s41582-023-00822-1.

5. Sofroniew, M. V., and Vinters, H. V. (2010). Astrocytes: Biology and pathology. Preprint, 10.1007/s00401-009-0619-8 10.1007/s00401-009-0619-8.

6. Anderson, M.A., Burda, J.E., Ren, Y., Ao, Y., O’Shea, T.M., Kawaguchi, R., Coppola, G., Khakh, B.S., Deming, T.J., and Sofroniew, M. V. (2016). Astrocyte scar formation AIDS central nervous system axon regeneration. Nature 532, 195–200. 10.1038/nature17623.

7. Pekny, M., and Pekna, M. (2014). ASTROCYTE REACTIVITY AND REACTIVE ASTROGLIOSIS: COSTS AND BENEFITS. Physiol Rev 94, 1077–1098. 10.1152/physrev.00041.2013.-Astrocytes.

8. Verkhratsky, A., Butt, A., Li, B., Illes, P., Zorec, R., Semyanov, A., Tang, Y., and Sofroniew, M. V. (2023). Astrocytes in human central nervous system diseases: a frontier for new therapies. Preprint at Springer Nature, 10.1038/s41392-023-01628-9 10.1038/s41392-023-01628-9.

9. Bardehle, S., Krüger, M., Buggenthin, F., Schwausch, J., Ninkovic, J., Clevers, H., Snippert, H.J., Theis, F.J., Meyer-Luehmann, M., Bechmann, I., et al. (2013). Live imaging of astrocyte responses to acute injury reveals selective juxtavascular proliferation. Nat Neurosci 16, 580–586. 10.1038/nn.3371.

10. Frik, J., Merl-Pham, J., Plesnila, N., Mattugini, N., Kjell, J., Kraska, J., Gómez, R.M., Hauck, S.M., Sirko, S., and Götz, M. (2018). Cross-talk between monocyte invasion and astrocyte proliferation regulates scarring in brain injury. EMBO Rep 19. 10.15252/embr.201745294.

11. Sirko, S., Behrendt, G., Johansson, P.A., Tripathi, P., Costa, M., Bek, S., Heinrich, C., Tiedt, S., Colak, D., Dichgans, M., et al. (2013). Reactive glia in the injured brain acquire stem cell properties in response to sonic hedgehog glia. Cell Stem Cell 12, 426–439. 10.1016/j.stem.2013.01.019.

12. Liddelow, S.A., Guttenplan, K.A., Clarke, L.E., Bennett, F.C., Bohlen, C.J., Schirmer, L., Bennett, M.L., Münch, A.E., Chung, W.S., Peterson, T.C., et al. (2017). Neurotoxic reactive astrocytes are induced by activated microglia. Nature 541, 481–487. 10.1038/nature21029.

13. Guttenplan, K.A., Weigel, M.K., Prakash, P., Wijewardhane, P.R., Hasel, P., Rufen-Blanchette, U., Münch, A.E., Blum, J.A., Fine, J., Neal, M.C., et al. (2021). Neurotoxic reactive astrocytes induce cell death via saturated lipids. Nature 599, 102–107. 10.1038/s41586-021-03960-y.

14. Guttenplan, K.A., Stafford, B.K., El-Danaf, R.N., Adler, D.I., Münch, A.E., Weigel, M.K., Huberman, A.D., and Liddelow, S.A. (2020). Neurotoxic Reactive Astrocytes Drive Neuronal Death after Retinal Injury. Cell Rep 31. 10.1016/j.celrep.2020.107776.

15. Conforti, P., Mezey, S., Nath, S., Chu, Y.H., Malik, S.C., Martínez Santamaría, J.C., Deshpande, S.S., Pous, L., Zieger, B., and Schachtrup, C. (2022). Fibrinogen regulates lesion border-forming reactive astrocyte properties after vascular damage. Glia 70, 1251–1266. 10.1002/glia.24166.

16. Nahon, D.M., Vila Cuenca, M., van den Hil, F.E., Hu, M., de Korte, T., Frimat, J.P., van den Maagdenberg, A.M.J.M., Mummery, C.L., and Orlova, V. V. (2024). Self-assembling 3D vessel-on-chip model with hiPSC-derived astrocytes. Stem Cell Reports 19, 946–956. 10.1016/j.stemcr.2024.05.006.

17. Campisi, M., Shin, Y., Osaki, T., Hajal, C., Chiono, V., and Kamm, R.D. (2018). 3D self-organized microvascular model of the human blood-brain barrier with endothelial cells, pericytes and astrocytes. Biomaterials 180, 117–129. 10.1016/j.biomaterials.2018.07.014.

18. Santos, R., Vadodaria, K.C., Jaeger, B.N., Mei, A., Lefcochilos-Fogelquist, S., Mendes, A.P.D., Erikson, G., Shokhirev, M., Randolph-Moore, L., Fredlender, C., et al. (2017). Differentiation of Inflammation-Responsive Astrocytes from Glial Progenitors Generated from Human Induced Pluripotent Stem Cells. Stem Cell Reports 8, 1757– 1769. 10.1016/j.stemcr.2017.05.011.

19. Sonsalla, G., Malpartida, A.B., Riedemann, T., Gusic, M., Rusha, E., Bulli, G., Najas, S., Janjic, A., Hersbach, B.A., Smialowski, P., et al. (2024). Direct neuronal reprogramming of NDUFS4 patient cells identifies the unfolded protein response as a novel general reprogramming hurdle. Neuron 112, 1117–1132.e9. 10.1016/j.neuron.2023.12.020.

20. Moeendarbary, E., Weber, I.P., Sheridan, G.K., Koser, D.E., Soleman, S., Haenzi, B., Bradbury, E.J., Fawcett, J., and Franze, K. (2017). The soft mechanical signature of glial scars in the central nervous system. Nat Commun 8. 10.1038/ncomms14787.

21. O’Shea, T.M., Ao, Y., Wang, S., Ren, Y., Cheng, A.L., Kawaguchi, R., Shi, Z., Swarup, V., and Sofroniew, M. V. (2024). Derivation and transcriptional reprogramming of border-forming wound repair astrocytes after spinal cord injury or stroke in mice. Nat Neurosci 27, 1505–1521. 10.1038/s41593-024-01684-6.

22. Magnusson, J.P., Zamboni, M., Santopolo, G., Mold, J.E., Barrientos-Somarribas, M., Talavera-Lopez, C., Andersson, B., and Frisén, J. (2020). Activation of a neural stem cell transcriptional program in parenchymal astrocytes. Elife 9, 1–25. 10.7554/ELIFE.59733.

23. Gascón, S., Murenu, E., Masserdotti, G., Ortega, F., Russo, G.L., Petrik, D., Deshpande, A., Heinrich, C., Karow, M., Robertson, S.P., et al. (2016). Identification and Successful Negotiation of a Metabolic Checkpoint in Direct Neuronal Reprogramming. Cell Stem Cell 18, 396–409. 10.1016/j.stem.2015.12.003.

24. Sirko, S., Schichor, C., Della Vecchia, P., Metzger, F., Sonsalla, G., Simon, T., Bürkle, M., Kalpazidou, S., Ninkovic, J., Masserdotti, G., et al. (2023). Injury-specific factors in the cerebrospinal fluid regulate astrocyte plasticity in the human brain. Nat Med 29, 3149–3161. 10.1038/s41591-023-02644-6.

25. Galarza, S., Crosby, A.J., Pak, C.H., and Peyton, S.R. (2020). Control of Astrocyte Quiescence and Activation in a Synthetic Brain Hydrogel. Adv Healthc Mater 9, 1–12. 10.1002/adhm.201901419.

26. Mosesson, M.W. (2003). Fibrinogen γ chain functions. Preprint, 10.1046/j.1538-7836.2003.00063.x 10.1046/j.1538-7836.2003.00063.x.

27. Gailit, J., Clarke, C., Newman, D., Tonnesen, M.G., Mosesson, M.W., and Clark, R.A.F. (1997). Human Fibroblasts Bind Directly to Fibrinogen at RGD Sites through Integrin alphavbeta3.

28. Thiagarajan, P., Rippon, A.J., and Farrell, D.H. (1996). Alternative Adhesion Sites in Human Fibrinogen for Vascular Endothelial Cells †.

29. Millesi, F., Mero, S., Semmler, L., Rad, A., Stadlmayr, S., Borger, A., Supper, P., Haertinger, M., Ploszczanski, L., Windberger, U., et al. (2023). Systematic Comparison of Commercial Hydrogels Revealed That a Synergy of Laminin and Strain-Stiffening Promotes Directed Migration of Neural Cells. ACS Appl Mater Interfaces 15, 12678–12695. 10.1021/acsami.2c20040.

30. Caliari, S.R., and Burdick, J. a (2016). A practical guide to hydrogels for cell culture. Nat Methods 13, 405–414. 10.1038/nmeth.3839.

31. Lattke, M., Goldstone, R., Ellis, J.K., Boeing, S., Jurado-Arjona, J., Marichal, N., MacRae, J.I., Berninger, B., and Guillemot, F. (2021). Extensive transcriptional and chromatin changes underlie astrocyte maturation in vivo and in culture. Nat Commun 12. 10.1038/s41467-021-24624-5.

32. Segel, M., Neumann, B., Hill, M.F.E., Weber, I.P., Viscomi, C., Zhao, C., Young, A., Agley, C.C., Thompson, A.J., Gonzalez, G.A., et al. (2019). Niche stiffness underlies the ageing of central nervous system progenitor cells. Nature 573, 130–134. 10.1038/s41586-019-1484-9.

33. Miroshnikova, Y.A., Mouw, J.K., Barnes, J.M., Pickup, M.W., Lakins, J.N., Kim, Y., Lobo, K., Persson, A.I., Reis, G.F., McKnigh, T.R., et al. (2016). Tissue mechanics promote IDH1-dependent HIF1α-tenascin C feedback to regulate glioblastoma aggression. Nat Cell Biol 18, 1336–1345. 10.1038/ncb3429.

34. Schmitt, A., Rödel, P., Anamur, C., Seeliger, C., Imhoff, A.B., Herbst, E., Vogt, S., Van Griensven, M., Winter, G., and Engert, J. (2015). Calcium alginate gels as stem cell matrix -making paracrine stem cell activity available for enhanced healing after surgery. PLoS One 10. 10.1371/journal.pone.0118937.

35. Matyash, M., Despang, F., Ikonomidou, C., and Gelinsky, M. (2014). Swelling and mechanical properties of alginate hydrogels with respect to promotion of neural growth. Tissue Eng Part C Methods 20, 401–411. 10.1089/ten.tec.2013.0252.

36. Tekin, H., Simmons, S., Cummings, B., Gao, L., Adiconis, X., Hession, C.C., Ghoshal, A., Dionne, D., Choudhury, S.R., Yesilyurt, V., et al. (2018). Effects of 3D culturing conditions on the transcriptomic profile of stem-cell-derived neurons. Nat Biomed Eng 2, 540–554. 10.1038/s41551-018-0219-9.

37. Salsmann, A., Schaffner-Reckinger, E., Kabile, F., Plançon, S., and Kieffer, N. (2005). A new functional role of the fibrinogen RGD motif as the molecular switch that selectively triggers integrin αIIbβ3-dependent RhoA activation during cell spreading. Journal of Biological Chemistry 280, 33610–33619. 10.1074/jbc.M500146200.

38. Gorodetsky, R., Vexler, A., Shamir, M., An, J., Levdansky, L., Shimeliovich, I., and Marx, G. (2003). New cell attachment peptide sequences from conserved epitopes in the carboxy termini of fibrinogen. Exp Cell Res 287, 116–129. 10.1016/S0014-4827(03)00120-4.

39. Reyhani, V., Seddigh, P., Guss, B., Gustafsson, R., Rask, L., and Rubin, K. (2014). Fibrin binds to collagen and provides a bridge for αVβ3 integrin-dependent contraction of collagen gels. Biochemical Journal 462, 113–123. 10.1042/BJ20140201.

40. Bocchi, R., Thorwirth, M., Simon-Ebert, T., Koupourtidou, C., Clavreul, S., Kolf, K., Della Vecchia, P., Bottes, S., Jessberger, S., Zhou, J., et al. (2025). Astrocyte heterogeneity reveals region-specific astrogenesis in the white matter. Nat Neurosci. 10.1038/s41593-025-01878-6.

41. Song, Y., Jiang, W., Afridi, S.K., Wang, T., Zhu, F., Xu, H., Nazir, F.H., Liu, C., Wang, Y., Long, Y., et al. (2024). Astrocyte-derived CHI3L1 signaling impairs neurogenesis and cognition in the demyelinated hippocampus. Cell Rep 43. 10.1016/j.celrep.2024.114226.

42. Zhang, Y., Sloan, S.A., Clarke, L.E., Caneda, C., Plaza, C.A., Blumenthal, P.D., Vogel, H., Steinberg, G.K., Edwards, M.S.B., Li, G., et al. (2016). Purification and Characterization of Progenitor and Mature Human Astrocytes Reveals Transcriptional and Functional Differences with Mouse. Neuron 89, 37–53. 10.1016/j.neuron.2015.11.013.

43. Santos, R., Vadodaria, K.C., Jaeger, B.N., Mei, A., Lefcochilos-Fogelquist, S., Mendes, A.P.D., Erikson, G., Shokhirev, M., Randolph-Moore, L., Fredlender, C., et al. (2017). Differentiation of Inflammation-Responsive Astrocytes from Glial Progenitors Generated from Human Induced Pluripotent Stem Cells. Stem Cell Reports 8, 1757– 1769. 10.1016/j.stemcr.2017.05.011.

44. Barbar, L., Jain, T., Zimmer, M., Kruglikov, I., Sadick, J.S., Wang, M., Kalpana, K., Rose, I.V.L., Burstein, S.R., Rusielewicz, T., et al. (2020). CD49f Is a Novel Marker of Functional and Reactive Human iPSC-Derived Astrocytes. Neuron 107, 436–453.e12. 10.1016/j.neuron.2020.05.014.

45. Sadick, J.S., O’Dea, M.R., Hasel, P., Dykstra, T., Faustin, A., and Liddelow, S.A. (2022). Astrocytes and oligodendrocytes undergo subtype-specific transcriptional changes in Alzheimer’s disease. Neuron 110, 1788–1805.e10. 10.1016/j.neuron.2022.03.008.

46. Bormann, D., Knoflach, M., Poreba, E., Riedl, C.J., Testa, G., Orset, C., Levilly, A., Cottereau, A., Jauk, P., Hametner, S., et al. (2024). Single-nucleus RNA sequencing reveals glial cell type-specific responses to ischemic stroke in male rodents. Nat Commun 15. 10.1038/s41467-024-50465-z.

47. Rempe, R.G., Hartz, A.M.S., and Bauer, B. (2016). Matrix metalloproteinases in the brain and blood-brain barrier: Versatile breakers and makers. Preprint at Nature Publishing Group, 10.1177/0271678X16655551 10.1177/0271678X16655551.

48. Liu, Y., Zhang, M., Hao, W., Mihaljevic, I., Liu, X., Xie, K., Walter, S., and Fassbender, K. (2013). Matrix metalloproteinase-12 contributes to neuroinflammation in the aged brain. Neurobiol Aging 34, 1231–1239. 10.1016/j.neurobiolaging.2012.10.015.

49. Wang, P., Gorter, R.P., de Jonge, J.C., Nazmuddin, M., Zhao, C., Amor, S., Hoekstra, D., and Baron, W. (2018). MMP7 cleaves remyelination-impairing fibronectin aggregates and its expression is reduced in chronic multiple sclerosis lesions. Glia 66, 1625–1643. 10.1002/glia.23328.

50. Man, J.H.K., Breur, M., van Gelder, C.A.G.H., Marcon, G., Maderna, E., Giaccone, G., Altelaar, M., van der Knaap, M.S., and Bugiani, M. (2024). Region-specific and age-related differences in astrocytes in the human brain. Neurobiol Aging 140, 102–115. 10.1016/j.neurobiolaging.2024.02.016.

51. Endo, F., Kasai, A., Soto, J.S., Yu, X., Qu, Z., Hashimoto, H., Gradinaru, V., Kawaguchi, R., and Khakh, B.S. (2022). Molecular basis of astrocyte diversity and morphology across the CNS in health and disease. Science (1979) 378. 10.1126/science.adc9020.

52. Robel, S., Bardehle, S., Lepier, A., Brakebusch, C., and Götz, M. (2011). Genetic deletion of Cdc42 reveals a crucial role for astrocyte recruitment to the injury site in vitro and in vivo. Journal of Neuroscience 31, 12471–12482. 10.1523/JNEUROSCI.2696-11.2011.

53. Puglisi, M., Lao, C.L., Wani, G., Masserdotti, G., Bocchi, R., and Götz, M. (2024). Comparing Viral Vectors and Fate Mapping Approaches for Astrocyte-to-Neuron Reprogramming in the Injured Mouse Cerebral Cortex. Cells 13. 10.3390/cells13171408.

54. Hand, R., Bortone, D., Mattar, P., Nguyen, L., Heng, J.I.T., Guerrier, S., Boutt, E., Peters, E., Barnes, A.P., Parras, C., et al. (2005). Phosphorylation of neurogenin2 specifies the migration properties and the dendritic morphology of pyramidal neurons in the neocortex. Neuron 48, 45–62. 10.1016/j.neuron.2005.08.032.

55. Ali, F., Hindley, C., McDowell, G., Deibler, R., Jones, A., Kirschner, M., Guillemot, F., and Philpott, A. (2011). Cell cycle-regulated multi-site phosphorylation of neurogenin 2 coordinates cell cycling with differentiation during neurogenesis. Development 138, 4267–4277. 10.1242/dev.067900.

56. Hindley, C., Ali, F., McDowell, G., Cheng, K., Jones, A., Guillemot, F., and Philpott, A. (2012). Post-translational modification of Ngn2 differentially affects transcription of distinct targets to regulate the balance between progenitor maintenance and differentiation. Development 139, 1718–1723. 10.1242/dev.077552.

57. Pereira, A., Diwakar, J., Masserdotti, G., Beşkardeş, S., Simon, T., So, Y., Martín-Loarte, L., Bergemann, F., Vasan, L., Schauer, T., et al. (2024). Direct neuronal reprogramming of mouse astrocytes is associated with multiscale epigenome remodeling and requires Yy1. Nat Neurosci 27, 1260–1273. 10.1038/s41593-024-01677-5.

58. Wang, M., Zhang, L., Novak, S.W., Yu, J., Gallina, I.S., Xu, L.L., Lim, C.K., Fernandes, S., Shokhirev, M.N., Williams, A.E., et al. (2024). Morphological diversification and functional maturation of human astrocytes in glia-enriched cortical organoid transplanted in mouse brain. Nat Biotechnol. 10.1038/s41587-024-02157-8.

59. Hajal, C., Offeddu, G.S., Shin, Y., Zhang, S., Morozova, O., Hickman, D., Knutson, C.G., and Kamm, R.D. (2022). Engineered human blood–brain barrier microfluidic model for vascular permeability analyses (Springer US) 10.1038/s41596-021-00635-w.

60. Yao, L., Sai, H.V., Shippy, T., and Li, B. (2024). Cellular and Transcriptional Response of Human Astrocytes to Hybrid Protein Materials. ACS Appl Bio Mater 7, 2887–2898. 10.1021/acsabm.3c01266.

61. Budday, S., Sommer, G., Birkl, C., Langkammer, C., Haybaeck, J., Kohnert, J., Bauer, M., Paulsen, F., Steinmann, P., Kuhl, E., et al. (2017). Mechanical characterization of human brain tissue. Acta Biomater 48, 319–340. 10.1016/j.actbio.2016.10.036.

62. Budday, S., Ovaert, T.C., Holzapfel, G.A., Steinmann, P., and Kuhl, E. (2019). Fifty Shades of Brain: A Review on the Mechanical Testing and Modeling of Brain Tissue (Springer Netherlands) 10.1007/s11831-019-09352-w.

63. Henrik Heiland, D., Ravi, V.M., Behringer, S.P., Frenking, J.H., Wurm, J., Joseph, K., Garrelfs, N.W.C., Strähle, J., Heynckes, S., Grauvogel, J., et al. (2019). Tumor-associated reactive astrocytes aid the evolution of immunosuppressive environment in glioblastoma. Nat Commun 10. 10.1038/s41467-019-10493-6.

64. Barbar, L., Rusielewicz, T., Zimmer, M., Kalpana, K., and Fossati, V. (2020). Isolation of Human CD49f+ Astrocytes and In Vitro iPSC-Based Neurotoxicity Assays. Neuron 1. 10.1016/j.xpro.2020.100172.

65. C. Benincasa, J., Madias, M.I., Kandell, R.M., Delgado-Garcia, L.M., Engler, A.J., Kwon, E.J., and Porcionatto, M.A. (2024). Mechanobiological Modulation of In Vitro Astrocyte Reactivity Using Variable Gel Stiffness. ACS Biomater Sci Eng 10, 4279– 4296. 10.1021/acsbiomaterials.4c00229.

66. Gomez-Cruz, C., Fernandez-de la Torre, M., Lachowski, D., Prados-de-Haro, M., del Río Hernández, A.E., Perea, G., Muñoz-Barrutia, A., and Garcia-Gonzalez, D. (2024). Mechanical and Functional Responses in Astrocytes under Alternating Deformation Modes Using Magneto-Active Substrates. Advanced Materials 36. 10.1002/adma.202312497.

67. Schachtrup, C., Ryu, J.K., Helmrick, M.J., Vagena, E., Galanakis, D.K., Degen, J.L., Margolis, R.U., and Akassoglou, K. (2010). Fibrinogen triggers astrocyte scar formation by promoting the availability of active TGF-β after vascular damage. Journal of Neuroscience 30, 5843–5854. 10.1523/JNEUROSCI.0137-10.2010.

68. Xiang, R., Wang, J., Chen, Z., Tao, J., Peng, Q., Ding, R., Zhou, T., Tu, Z., Wang, S., Yang, T., et al. (2025). Spatiotemporal transcriptomic maps of mouse intracerebral hemorrhage at single-cell resolution. Neuron. 10.1016/j.neuron.2025.04.026.

69. Kajtez, J., Laurin, K., Nilsson, F., Bruzelius, A., Cepeda-Prado, E., Birtele, M., Barker, R.A., Herborg, F., Rylander Ottosson, D., Storm, P., et al. (2025). Three-dimensional cell-cell interactions promote direct reprogramming of patient fibroblasts into functional and transplantable neurons.

70. Jin, Y., Lee, J.S., Kim, J., Min, S., Wi, S., Yu, J.H., Chang, G.-E., Cho, A.-N., Choi, Y., Ahn, D.-H., et al. (2018). Three-dimensional brain-like microenvironments facilitate the direct reprogramming of fibroblasts into therapeutic neurons. Nat Biomed Eng 2, 522–539. 10.1038/s41551-018-0260-8.

71. Sun, Z., Kwon, J.S., Ren, Y., Chen, S., Walker, C.K., Lu, X., Cates, K., Karahan, H., Sviben, S., Fitzpatrick, J.A.J., et al. (2024). Modeling late-onset Alzheimer’s disease neuropathology via direct neuronal reprogramming. Science 385, adl2992. 10.1126/science.adl2992.

72. Götz, M., Sirko, S., Beckers, J., and Irmler, M. (2015). Reactive astrocytes as neural stem or progenitor cells: In vivo lineage, In vitro potential, and Genome-wide expression analysis. Preprint at John Wiley and Sons Inc., 10.1002/glia.22850 10.1002/glia.22850.

73. Chen, W., Guillaume-Gentil, O., Rainer, P.Y., Gäbelein, C.G., Saelens, W., Gardeux, V., Klaeger, A., Dainese, R., Zachara, M., Zambelli, T., et al. (2022). Live-seq enables temporal transcriptomic recording of single cells. Nature 608, 733–740. 10.1038/s41586-022-05046-9.

74. Sohrabi, A., Lefebvre, A.E.Y.T., Harrison, M.J., Condro, M.C., Sanazzaro, T.M., Safarians, G., Solomon, I., Bastola, S., Kordbacheh, S., Toh, N., et al. (2023). Microenvironmental stiffness induces metabolic reprogramming in glioblastoma. Cell Rep 42. 10.1016/j.celrep.2023.113175.

75. Campisi, M., Shin, Y., Osaki, T., Hajal, C., Chiono, V., and Kamm, R.D. (2018). 3D self-organized microvascular model of the human blood-brain barrier with endothelial cells, pericytes and astrocytes. Biomaterials 180, 117–129. 10.1016/j.biomaterials.2018.07.014.

76. LeSavage, B.L., Suhar, N.A., Madl, C.M., and Heilshorn, S.C. (2018). Production of Elastin-like Protein Hydrogels for Encapsulation and Immunostaining of Cells in 3D. Journal of Visualized Experiments. 10.3791/57739-v.

77. Vorwald, C.E., Gonzalez-Fernandez, T., Joshee, S., Sikorski, P., and Leach, J.K. (2020). Tunable fibrin-alginate interpenetrating network hydrogels to support cell spreading and network formation. Acta Biomater 108, 142–152. 10.1016/j.actbio.2020.03.014.

78. Hutter, J.L., and Bechhoefer, J. (1993). Calibration of atomic-force microscope tips. Review of Scientific Instruments 64, 1868–1873. 10.1063/1.1143970.

79. Schindelin, J., Arganda-Carreras, I., Frise, E., Kaynig, V., Longair, M., Pietzsch, T., Preibisch, S., Rueden, C., Saalfeld, S., Schmid, B., et al. (2012). Fiji: An open-source platform for biological-image analysis. Preprint, 10.1038/nmeth.2019 10.1038/nmeth.2019.

80. Feng, L., Zhao, T., and Kim, J. (2015). Neutube 1.0: A new design for efficient neuron reconstruction software based on the swc format. eNeuro 2. 10.1523/ENEURO.0049-14.2014.

81. Arshadi, C., Günther, U., Eddison, M., Harrington, K.I.S., and Ferreira, T.A. (2021). SNT: a unifying toolbox for quantification of neuronal anatomy. Nat Methods 18, 374–377. 10.1038/s41592-021-01105-7.

82. Bolger, A.M., Lohse, M., and Usadel, B. (2014). Trimmomatic: A flexible trimmer for Illumina sequence data. Bioinformatics 30, 2114–2120. 10.1093/bioinformatics/btu170. https://netcologne.dl.sourceforge.net/project/rseqc/BED/Human_Homo_sapiens/hg38_Gencode_V28.bed.gz.

83. Alexa, A., and Rahnenfuhrer, J. (2024). topGO: Enrichment Analysis for Gene Ontology. Bioconductor. 10.18129/B9.bioc.topGO.

84. Heinrich, C., Blum, R., Gascón, S., Masserdotti, G., Tripathi, P., Sánchez, R., Tiedt, S., Schroeder, T., Götz, M., and Berninger, B. (2010). Directing astroglia from the cerebral cortex into subtype specific functional neurons. PLoS Biol 8. 10.1371/journal.pbio.1000373.

85. Heinrich, C., Gascón, S., Masserdotti, G., Lepier, A., Sanchez, R., Simon-Ebert, T., Schroeder, T., G’tz, M., and Berninger, B. (2011). Generation of subtype-specific neurons from postnatal astroglia of the mouse cerebral cortex. Nat Protoc 6, 214–228. 10.1038/nprot.2010.188.

86. Drouin-Ouellet, J., Lau, S., Brattås, P.L., Rylander Ottosson, D., Pircs, K., Grassi, D.A., Collins, L.M., Vuono, R., Andersson Sjöland, A., Westergren-Thorsson, G., et al. (2017). REST suppression mediates neural conversion of adult human fibroblasts via microRNA-dependent and -independent pathways. EMBO Mol Med 9, 1117–1131. 10.15252/emmm.201607471.

87. Love, M.I., Huber, W., and Anders, S. (2014). Moderated estimation of fold change and dispersion for RNA-seq data with DESeq2. Genome Biol 15, 550. 10.1186/s13059-014-0550-8.

88. Hadley Wickham (2016). ggplot2 - Elegant Graphics for Data Analysis (2nd Edition) (Springer-Verlag).

