## Supplementary material for "Fibrin-containing Hydrogels Regulate Human Astrocyte State and Neuronal Reprogramming": Figures S

### Supplementary Figures

#### Figure S1

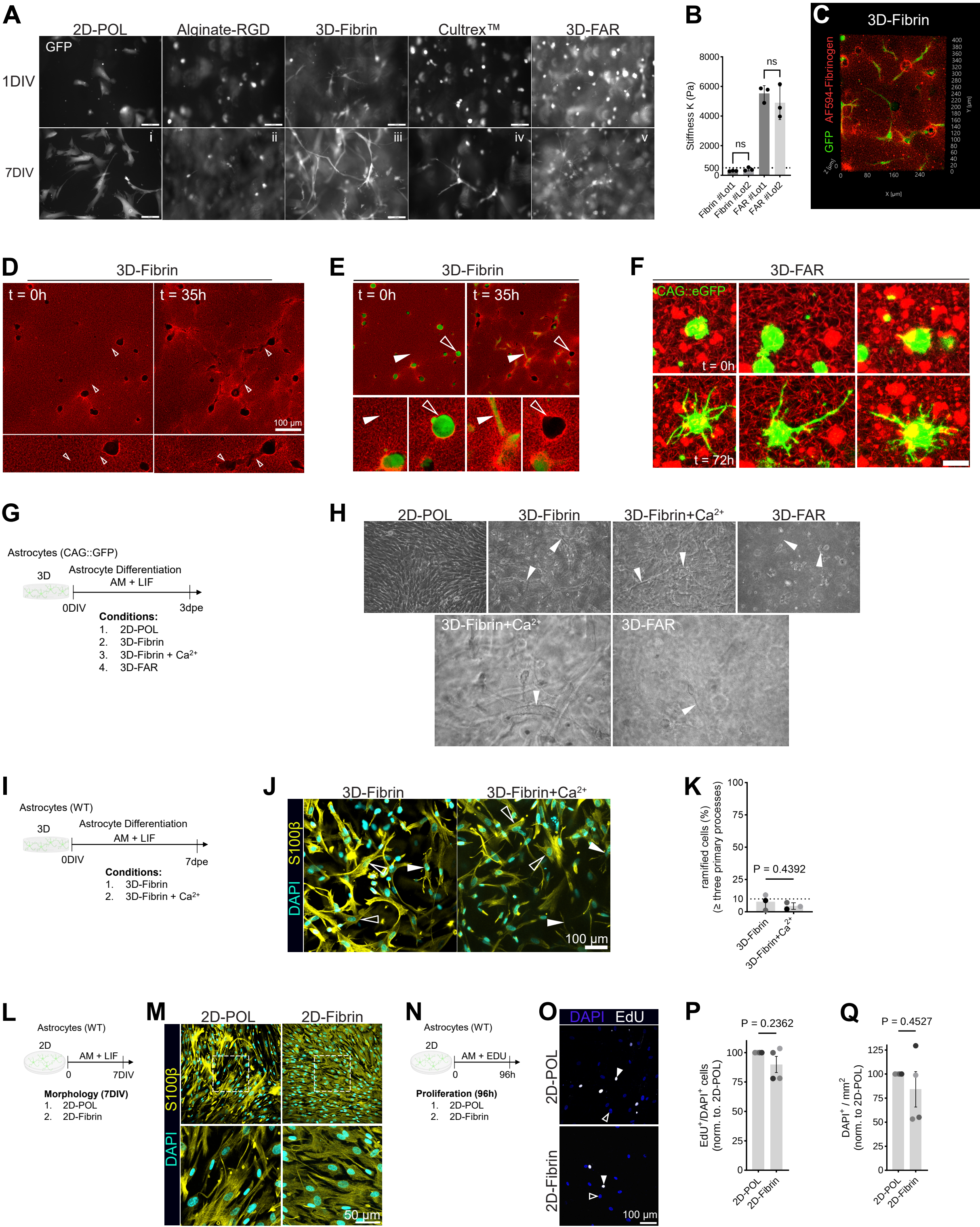

**Figure S1 – hiPSC-derived astrocytes grown in different 3D hydrogel substrates and LIVE-cell imaging**

(**A**) Epifluorescence microscopy images of $GFP^{+}$ astrocytes embedded in different 3D hydrogel materials for seven days *in vitro*. Scale bars = 100 µm. (**B**) Stiffness of 3D-Fibrin and 3D-FAR hydrogels prepared from two different batches of fibrinogen. (N = 3, mean ± SD, one-way ANOVA test, ns: $p \geq 0.05$). (**C**) Example of a astrocyte (CAG::GFP pAstro) inside 3D-Fibrin hydrogel (Alexa Fluor 594-conjugated, red). (**D**) Pictures showing human astrocytes embedded in 3D-Fibrin at 0 and at 35 hours. Scalebar = 100µm. (**E**) Pcitures depicting astrocyte migration inside 3D-Fibrin hydrogels over time. (**F**) Example of morphological changes of an astrocyte embedded in 3D-FAR hydogel over time. Scale bar = 20 µm. (**G**) Experimental design to test the effect of $Ca^{2+}$ exposure on astrocytes in 2D and in 3D at 3 dpe. (**H**) Brightfield images revealing the morphology of astrocytes embedded in Fibrin or 3D-Fibrin treated with $Ca^{2+}$ (white arrows), compared to 3D-FAR embedded astrocytes (yellow arrows). (**I**) Experimental design to test the effect of $Ca^{2+}$ exposure on astrocytes in 2D and in 3D at 7 dpe. (**J**) Pictures showing $S100\beta^{+}DAPI^{+}$ astrocytes in 3D-Fibrin or 3D-Fibrin+$Ca^{2+}$, with ramified cells (white arrows) and not-ramified cells (unfilled white arrows) marked. **(K)** Graph depicting the proportion of ramified cells (≥ three main primary processes) in 3D-Fibrin or 3D-Fibrin+$Ca^{2+}$ (N = 3, mean ± SD, paired student's t-test). (**L**) Experimental design to assess the morphology of astrocytes grown on 2D-POL or 2D-Fibrin hydrogels for 7 DIV. (**M**) Pictures of $S100\beta^{+}DAPI^{+}$ astrocytes on 2D-POL or 2D-Fibrin. (**N**) Experimental design of assessing proliferation of astrocytes either cultured for 96h on 2D-POL or on 2D-Fibrin by EdU administration. (**O**) Examples of cells on 2D-POL or 2D-Fibrin positive for EdU. Quantification of $EdU^{+}/DAPI^{+}$ cells, normalized to the amount of $EdU^{+}$ astrocytes (white arrows) on 2D-POL (glass) (N = 4, mean ± S.E.M). (**P-Q**) Graphs showing the proportion of $EdU^{+}/DAPI^{+}$ cells, normalized to the amount of $EdU^{+}$ astrocytes (white arrows) on 2D-POL (N = 4, mean ± S.E.M, paired student's t-test, ) and cell density ($DAPI^{+}$ cells per $mm^2$ hydrogel or POL-surface) after 96 h on 2D-POL or 2D-Fibrin, normalized to the amount of cells on 2D-POL (N = 4, mean ± S.E.M, paired student's t-test).

#### Figure S2

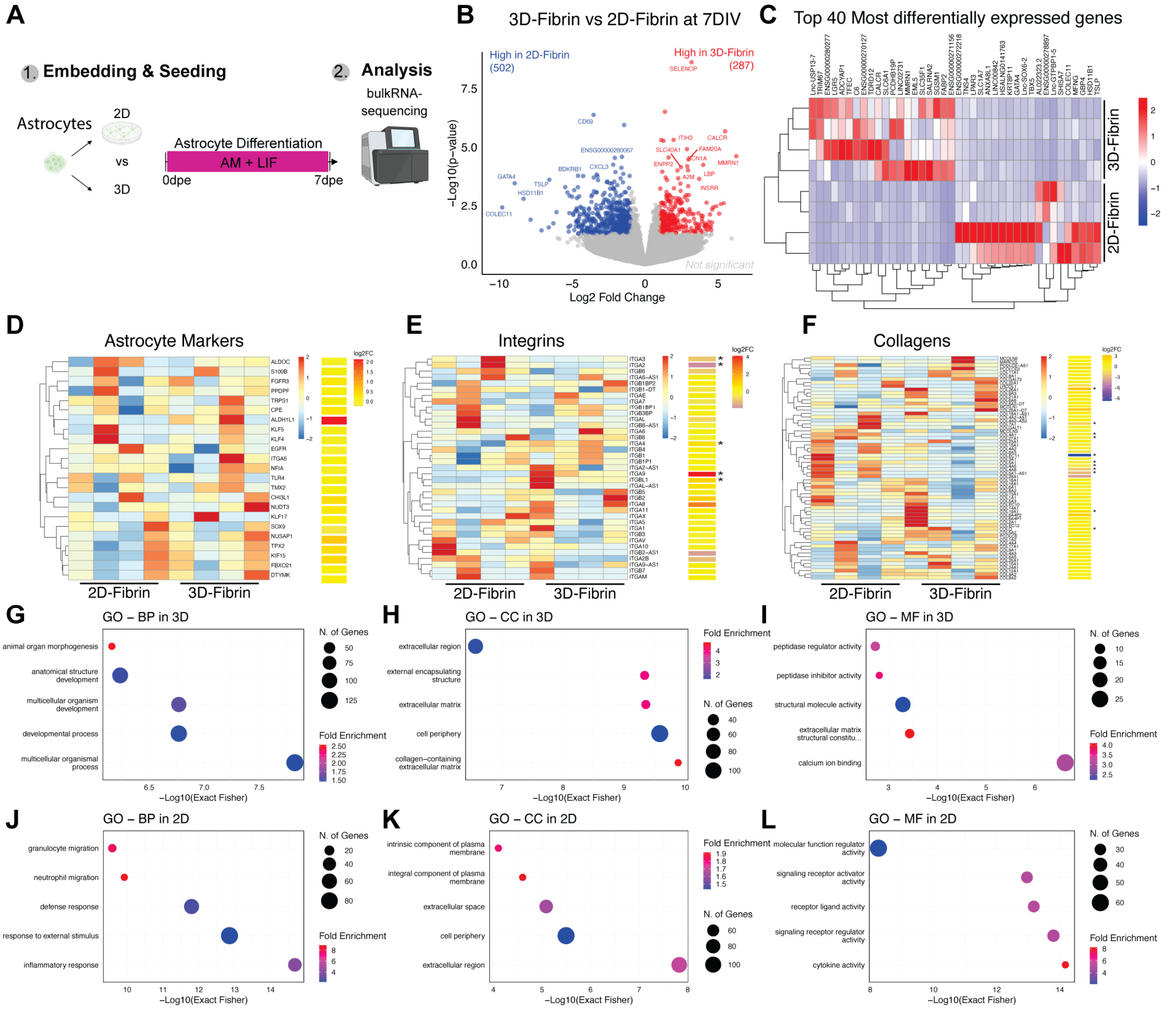


**Figure S2 – Bulk RNA-sequencing of astrocytes on 2D-Fibrin and in 3D-Fibrin**

(**A**) Experimental design: astrocytes, cultured on 2D-Fibrin or 3D-Fibrin, were collected at 7 dpe and analyzed via RNA-sequencing. (**B**) Volcano plot showing differentially expressed genes in astrocyte cultured in 3D-Fibrin (red; log2FC>1, pvalue<0.05) or in 2D-Fibrin (blue; log2FC< -1, pvalue<0.05). Names are indicated when log2FC> abs(2). (**C**) Heatmap showing the top 40 genes more expressed by astrocytes in 3D-Fibrin or 2D-Fibrin. (**D-F**) Heatmaps (left panels) showing the relative expression of astrocyte markers (D), integrins (E), collagens (F). In each panel, the right heatmap depicts the log2fold change according to RNA-seq analysis. Asterisks indicate their statistically significance (*pval<0.05). (**G-I**) Gene ontology (GO) analysis of biological processes (BP, G), cellular compartment (CC, H) and molecular function (MF, I) from genes more expressed in 3D-Fibrin (log2FC>1, pval<0.05). (**J-L**) GO analysis of biological processes (BP, J), cellular compartment (CC, K) and molecular function (MF, L) from genes more expressed in 2D-Fibrin (log2FC< -1, pval<0.05).

### Figure S3

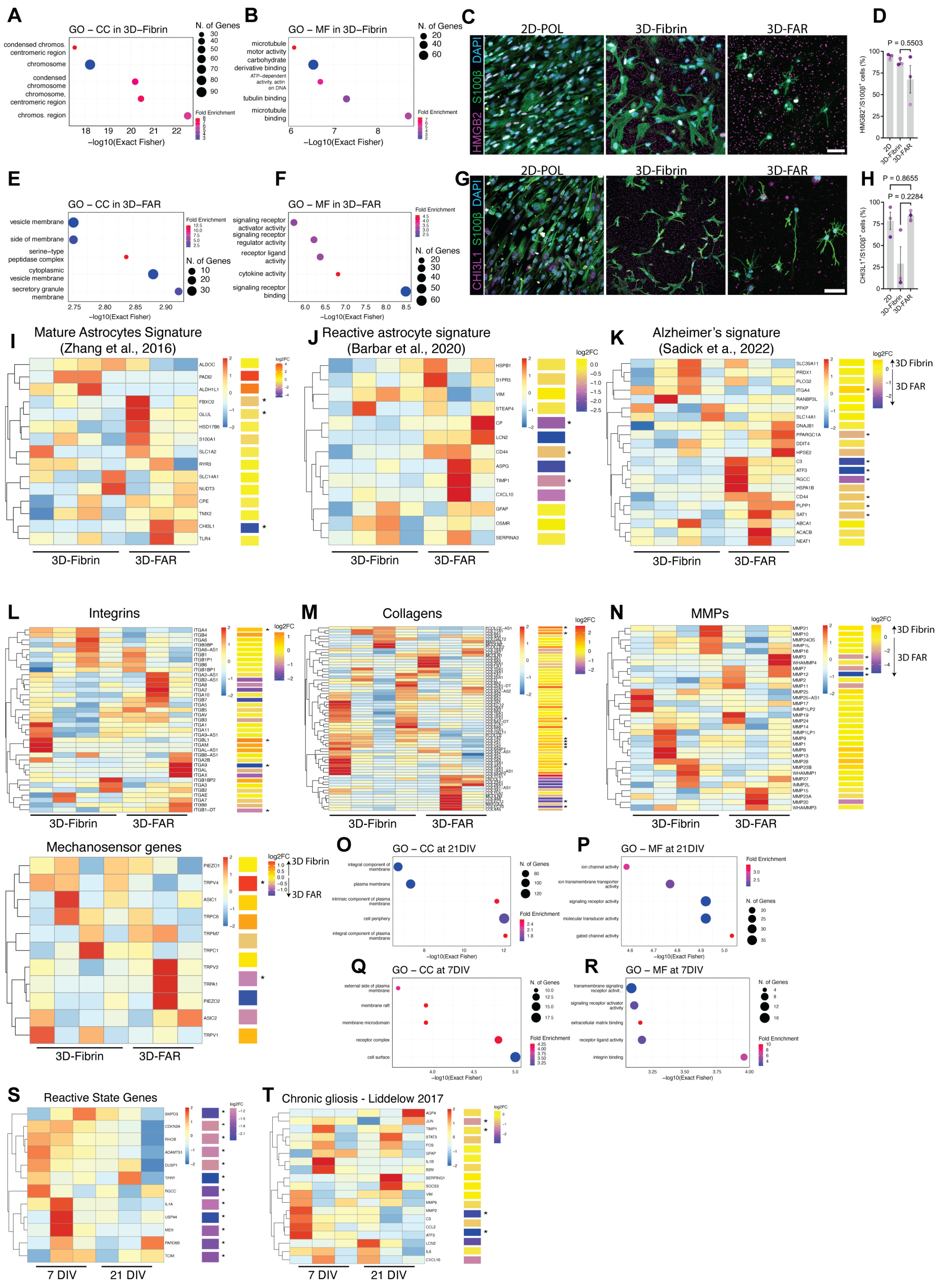

A
GO – CC in 3D–Fibrin
B
GO – MF in 3D–Fibrin
C
2D-POL
3D-Fibrin
3D-FAR
D
P = 0.5503
E
GO – CC in 3D–FAR
F
GO – MF in 3D–FAR
G
H
P = 0.8655
P = 0.2284
I
Mature Astrocytes Signature (Zhang et al., 2016)
J
Reactive astrocyte signature (Barbar et al., 2020)
K
Alzheimer's signature (Sadick et a., 2022)
L
Integrins
M
Collagens
N
MMPs
Mechanosensor genes
O
GO – CC at 21DIV
P
GO – MF at 21DIV
Q
GO – CC at 7DIV
R
GO – MF at 7DIV
S
Reactive State Genes
T
Chronic gliosis - Liddelow 2017
3D-Fibrin
3D-FAR
7 DIV
21 DIV

**Figure S3 – Bulk-RNA seq analysis and validation**

(**A, B, E, F**) GO analysis of cellular compartment (CC, A, E) and molecular function (MF, B, F) from genes more expressed in 3D-Fibrin (log2FC>1, pval<0.05, A, B) or in 3D-FAR (log2FC>1, pval<0.05, E, F). (**C, D**) Pictures showing the expression of HMGB2 in astrocytes in different substrates at 7 dpe (C) and graph showing the proportion of astrocytes ($S100\beta^{+}$) positive for HMGB2 (D) (N = 3, mean ± S.E.M). (**G, H**) Pictures showing the expression of CHI3L1 in astrocytes in different substrates at 7 dpe (G) and graph showing the proportion of astrocytes ($S100\beta^{+}$) positive for CHI3L1 (D) (N = 3, mean ± S.E.M). (**I-O**) Heatmaps (left panels) showing the relative expression of genes associated to mature astrocytes (I), reactive gliosis (J), Alzheimer's disease(K), genes coding integrins (L), collagens (M), metalloproteases (MMPs, N) and mechanosensitive proteins (O) in 3D-Fibrin and 3D-FAR at 7 dpe. In each panel, the right heatmap depicts the log2fold change according to RNA-seq analysis. Asterisks indicate their statistically significance (*pval<0.05). (**P-S**) GO analysis of cellular compartment (CC, P, R) and molecular function (MF, Q, S) from genes more expressed in 3D-FAR at 21 dpe (log2FC>1, pval<0.05, N, O) or at 21 dpe (log2FC>1, pval<0.05, P, Q). (**T, U**) Heatmaps (left panels) showing the relative expression of genes associated to inflammatory signature (from GO terms of the GO term analysis of 3D-FAR vs 3D-Fibrin) (T) and chronic gliosis (U) (from Liddelow et al. 2017) in 3D-FAR at 7 and 21 dpe. In each panel, the right heatmap depicts the log2fold change according to RNA-seq analysis. Asterisks indicate statistically significant changes between 7 and 21 dpe expression (*pval<0.05).

### Figure S4

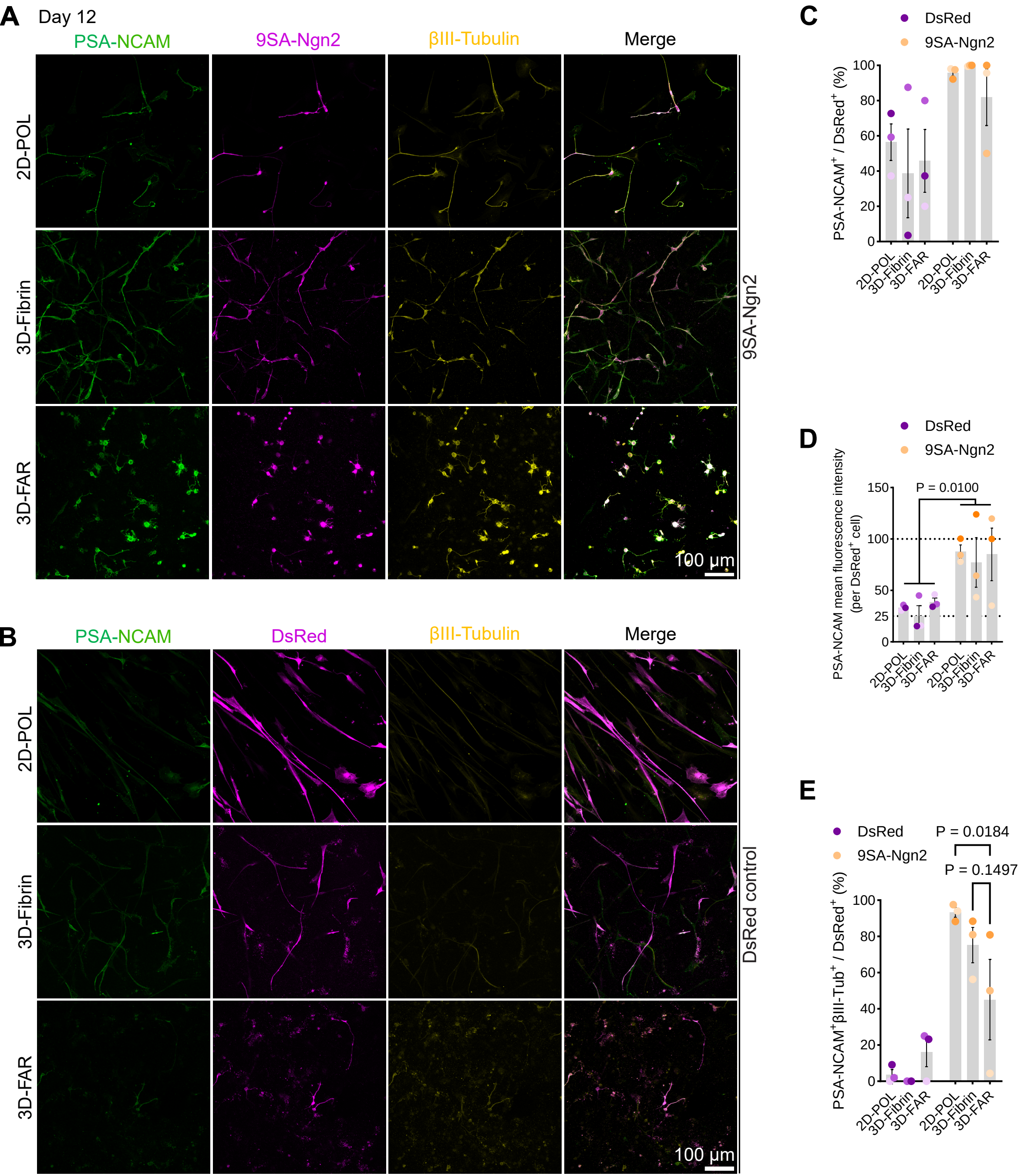
A
Day 12
PSA-NCAM
9SA-Ngn2
βIII-Tubulin
Merge
2D-POL
3D-Fibrin
3D-FAR
9SA-Ngn2
100 µm
B
PSA-NCAM
DsRed
βIII-Tubulin
Merge
2D-POL
3D-Fibrin
3D-FAR
DsRed control
100 µm
C
DsRed
9SA-Ngn2
PSA-NCAM+ / DsRed+ (%)
2D-POL
3D-Fibrin
3D-FAR
D
DsRed
9SA-Ngn2
P = 0.0100
PSA-NCAM mean fluorescence intensity (per DsRed+ cell)
E
DsRed
9SA-Ngn2
P = 0.0184
P = 0.1497
PSA-NCAM+βIII-Tub+ / DsRed+ (%)

**Figure S4 – Early conversion of astrocytes after 12 dpe**

(**A, B**) Examples of DsRed$^+$PSA-NCAM$^+$βIII-Tubulin$^+$ astrocytes 12 dpe (**A**, RV-9SA-Ngn2. **B**, RV-DsRed control). Scalebar = 100μm. (**C**) Graph showing the proportion of PSA-NCAM$^+$/DsRed$^+$ cells (N = 3, mean ± S.E.M, colours indicate data from independent biological experiments). (**D**) Graph showing the PSA-NCAM fluorescence intensity among DsRed$^+$ cells in different conditions at 12 dpe (N = 3, mean ± S.E.M, colours indicate data from independent biological experiments). (**E**) Graph showing the conversion rate (NCAM$^+$βIII-Tubulin$^+$-DsRed$^+$ over DsRed$^+$ cells) at 12 dpe (N = 3, mean ± S.E.M, colours indicate data from independent biological experiments).
